# Persistent large-scale changes in alternative splicing in prefrontal cortical neuron types following psychedelic exposure

**DOI:** 10.1101/2025.01.16.633439

**Authors:** Yungyi Hsiao, Marcos AS Fonseca, Alina S Tiemroth, Estefany J Vasquez, Andrea M Gomez

## Abstract

Psychedelics engage the serotonergic system as potent neuromodulators, increasing neuroplasticity in humans and rodents. Persistent changes in cognitive flexibility, emotional regulation, and social cognition are thought to underlie the therapeutic effects of psychedelics. However, the underlying molecular and cellular basis of psychedelic-induced plasticity remains unclear. Here, we identify persistent, cell type-specific alternative splicing changes in the mouse medial prefrontal cortex (mPFC) induced by a single dose of psychedelics. Combining deep RiboTag sequencing and bioinformatics, we find that a single dose of psychedelics modestly alters gene expression while dramatically shifting patterns of alternative splicing lasting at least a month. We connect our functional enrichment and alternative splicing analysis with changes in the extracellular matrix, synaptic physiology, and intrinsic physiology in parvalbumin interneurons days to a week after psychedelic treatment. Our dataset is an essential resource for understanding the persistent, cell type-specific effects of psychedelics on cortical cell types and functions.

## Introduction

In Phase 2 double-blind, randomized clinical trials, a single dose of a psychedelic, such as psilocybin, provides fast-acting and persistent relief from neuropsychiatric disorders such as treatment-resistant depression, major depressive disorder, and cancer-related distress^1–4^. Recent efforts to understand the molecular and cellular basis by which psychedelics promote functional and behavioral changes in rodents yield mixed results regarding which cell types mediate the effects, the downstream molecular pathways involved, and how a single dose of psychedelics can lead to lasting changes in the brain ^5–10^.

Psychedelic-induced cellular and molecular changes have been found in brain regions relevant to behaviors in cognitive flexibility and emotional regulation, such as the cortex (somatosensory, visual, frontal cortices), amygdala, striatum, and hippocampus ^5–9,11–14^. Structurally similar to the neuromodulator serotonin, psychedelics engage the serotonergic system and induce neural plasticity in the brain ^15–17^. Classic psychedelics are divided into two main classes based on their chemical structure: tryptamines (e.g., psilocybin, N,N-dimethyltryptamine [DMT], and the ergoline, lysergic acid diethylamide [LSD]) and phenethylamines (e.g., mescaline, 2,5-Dimethoxy-4-iodoamphetamine [DOI]) ^18^. While classic psychedelics exhibit polypharmacology by engaging various neuromodulator receptors, signal transduction via the serotonin 2A receptor (5HT_2A_R) is critical for their psychoactivity ^11,19–24^. It remains unclear, however, whether tryptamine or phenethylamine psychedelics classes engage overlapping or distinct long-term molecular responses.

Our study focuses on the medial prefrontal cortex (mPFC), a brain region abundant in 5HT_2A_R in both mice and humans ^25,26^. The mPFC is central to the pathophysiology of neuropsychiatric disorders such as stress, depression, post-traumatic stress disorder, and schizophrenia ^27–31^. Although psychedelics alter excitatory synaptic transmission in the PFC ^5,32^, emerging evidence suggests that psychedelics may differentially affect excitatory and inhibitory cell types ^6–8^. Inhibition strongly regulates cortical plasticity and function, yet how inhibitory interneurons like parvalbumin (PV) interneurons contribute to the long-term psychedelic response remains unclear ^7,33–36^. In this study, we take a cell type-specific approach to understand the differential effects of psychedelics on key inhibitory and excitatory cortical cell types ^26^.

At the cellular level, psychedelics induce structural and functional plasticity that persists for up to a month after exposure in mice ^5,32,37,38^. Efforts to identify the underlying mechanism of psychedelic-induced plasticity have implicated differential gene expression as the basis for long-term plasticity ^5–7^. In addition to the acute induction of immediate early genes (IEGs) ^11,20^, recent transcriptomic studies identified changes in the expression of genes that control axon growth, synaptic transmission, and cell-cell adhesion, lasting days after exposure ^5–7,9^. However, it is not known if transcriptomic changes persist at longer time points. Moreover, previous studies using single-nuclei or single-cell RNA-seq (snRNA-seq or scRNA-seq), methods that remove all dendritic– and axon-localized RNAs, lacked sufficient read depth to quantify RNA isoforms, thereby overlooking the non-transcriptional roles, including changes in alternative RNA splicing, in the response to psychedelic exposure ^6–8,10^.

Activity-dependent transcription is necessary for the synthesis of new RNAs during neural plasticity, whereas non-transcriptional processes such as alternative splicing, translation, and post-translational modifications are critical for regulating mRNA stability, transport, and translation efficiency in order to adjust the levels and function of proteins ^39,40^. RNA splicing is a critical intermediate step between transcription and translation. Combinatorial use of alternative exons gives rise to numerous RNA isoforms, expanding protein diversity and function ^41^. The nervous system relies extensively upon alternative splicing to expand the coding capacity of the genome, giving rise to phenotypic diversity and is critical for establishing the synaptic properties of neurons ^42–44^. Further, alternative splicing exhibits greater cell type-specific patterns than expression in genes encoding proteins critical for the intrinsic and synaptic properties of neurons ^45^. Alternative splicing is dynamic, undergoing rapid changes during neuronal activity and drug exposure, and is altered in disease states ^39,46–49^ thus raising the possibility that changes in alternative splicing play a key role in psychedelic-induced plasticity.

Here, we present a dataset of high-depth, cell type-specific, longitudinal gene regulatory changes associated with a single dose of DOI or psilocybin across persistent time points with saline-matched controls. We identify modest changes in gene expression that return to baseline by a month, with PV interneurons exhibiting the greatest magnitude of change. Remarkably, despite a lack of gene expression changes, all neurons exhibit large degrees of alternative splicing changes that are selective to cell type and persistent for at least one month. Unlike prior studies, which focus on transcriptional roles of psychedelic-induced plasticity, our study suggests that non-transcriptional roles may be critical for the long-term effects of psychedelics. We connect the changes in gene regulation to functional changes by combining our RNA-seq data with histology and electrophysiology. Together, our dataset provides an essential resource for understanding the various cell type-specific effects of psychedelics on cortical cell types and functions.

## Results

### Deep sequencing of ribosome-bound mRNAs following psychedelic exposure in the mPFC

Though the mPFC comprises diverse excitatory and inhibitory neurons with distinct laminar positions and functions, we lack an understanding of the sustained effects of psychedelics across these cell populations ^50,51^. To better understand the molecular basis of persistent psychedelic-induced plasticity within cortical microcircuits, we used tagged-ribosome affinity purification (RiboTag) and performed deep sequencing of cell type-specific mRNAs from the adult (8-11 weeks) mouse mPFC at multiple extended time points (48 hours, 1 week, 1 month) following psychedelic treatment (Figure 1A) ^52^. Using the RiboTag approach, we gain cell type-specificity, preserve the detection of lowly expressed genes, and obtain deep coverage of splice junctions that is not achievable with snRNA-seq or scRNA-seq that sequences only the extreme 3’ end of the mRNA ^45^. Importantly, we focus on the translatome instead of the transcriptome by sequencing ribosome-bound mRNAs, while capturing RNAs localized throughout the cell body, including axons and dendrites ^53,54^.

**Figure 1.**
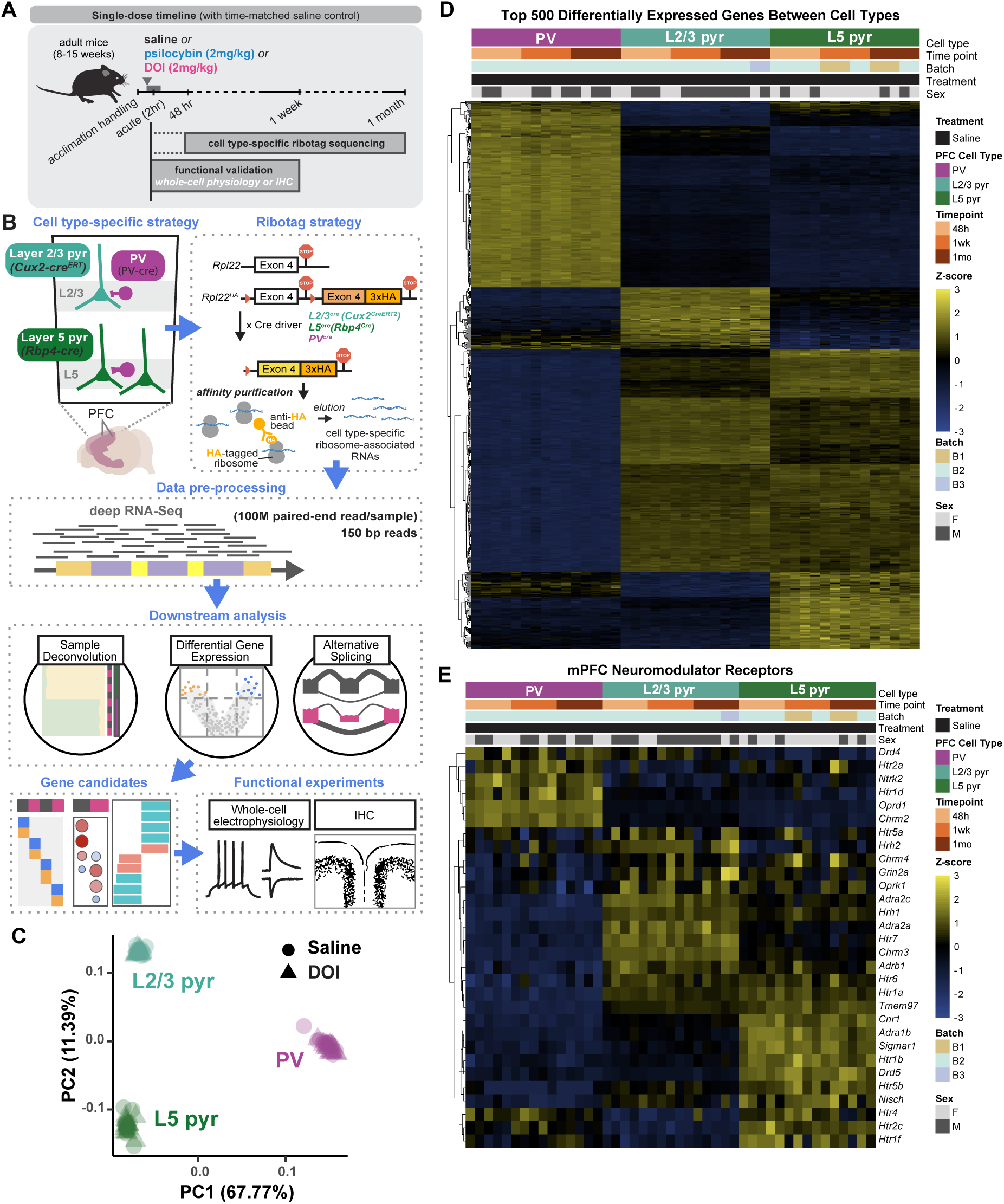
Cell type-specific, deep sequencing of ribosome-bound mRNAs following psychedelic exposure in the prefrontal cortex. **(A)** Timeline of translatomic and functional experiments in adult mice (8-15 weeks) at two-hours, 48-hours, 1-week, and 1-month following injection of DOI (2mg/kg), psilocybin (2mg/kg), or saline-matched controls. **(B)** Schematic of RiboTag experiments with downstream data processing and analysis, followed by functional characterization of phenotypes. **(C)** Principal component analysis (PCA) of all saline– and DOI-treated samples (n=90). **(D)** Heatmap showing expression of top 500 differentially expressed genes (DEGs) between cell types in all saline samples (|log_2_FC| > 1 and q < .05, top 200 genes from L2/3 vs. L5, top 300 unique genes from PV vs. both excitatory cell types), normalized by variance stabilizing transformation. **(E)** Heatmap showing cell type-specific expression of 30 neuromodulator receptors in the mPFC in all saline samples. See also Figure S1.

Conditional tagging of the ribosomal protein Rpl22 with hemagglutinin (HA) using Cre drivers allows for the purification of cell type-specific mRNAs ^52^. We generated mice that were homozygous for the *Rpl22-HA (RiboHA)* allele and heterozygous for the Cre alleles *PV-cre, Cux2^ERT^-cre*, or *Rbp4-cre* to label parvalbumin (PV) interneurons, layer 2/3 pyramidal neurons (L2/3), and layer 5 (L5) pyramidal neurons, respectively (Figure 1B) ^55^. Our approach allows us to compare the effects of psychedelics on excitatory and inhibitory neurons, as well as excitatory neurons with different laminar positions. We confirmed the specificity of the ribosome labeling in the mPFC with the Cre drivers using immunohistochemistry and RT-qPCR, which show enrichment and depletion of cell type-specific markers in target cell types (Figures S1A-S1C). We collected the mPFC 48 hours, one week, or up to one month following treatment with DOI (2mg/kg, i.p.) or saline-matched controls (n = 5 animals per cell type, per time point, and per treatment). To compare the responses of the phenylethylamine, DOI, with the tryptamine class, we also collected mPFC 48 hours following treatment with psilocybin (2mg/kg, i.p.).

Deep sequencing of the mRNAs (>100 million paired-end reads per sample, 150 base pairs read length, an average of 19,000 genes detected) resulted in an overall alignment rate of 79% (Table S1). Principal component analysis (PCA) comparing DOI and saline treatments (n = 90) confirmed that the primary source of variance between samples is cell type, affirming that neither psychedelic treatment nor time point affects cell identity (Figure 1C). To validate the quality of our dataset and verify cell type differences, differential gene expression analysis confirmed significant differences between PV interneurons and the two pyramidal neurons cell types, with fewer but notable differences between the L2/3 and L5 pyramidal neurons, and appropriate enrichment and de-enrichment of known cell type-specific markers (Figures 1D and Table S2).

Given the various affinities of psychedelics to multiple receptors in the brain, we assessed the distribution of neuromodulator receptors across our target PFC cell types, such as the different serotonin receptor subtypes (Figures 1E and S1D). We observed expression of 5HT_2A_R across all three cell types, with modest enrichment in L5 relative to L2/3 pyramidal neurons (4.84 FPKM vs. 3.98 FPKM) ^26^. Additionally, we noted the robust expression of 5HT_2A_R in PV interneurons (5.72 FPKM), suggesting a prominent role of PV interneurons in responding to psychedelics. We also observed differences in the expression of other serotonin receptors between cell types, such as 5HT_1A_R and 5HT_7_R, which may impact cell type-specific responses to psychedelic activity (Figure S1D). Our dataset thus provides a comprehensive profile of gene expression in the mPFC.

To further characterize the cell type compositions represented in our dataset based on established transcriptional signatures, we applied MUlti-Subject SIngle Cell (MuSiC) deconvolution trained on scRNA-seq atlas from the mouse visual cortex ^56,57^. MuSiC deconvolution of our RiboTag samples revealed an overwhelming majority of excitatory cell type in our L2/3 and L5 pyramidal neuron samples and strictly interneurons in the PV samples, with little to no representation of non-neuronal cell types (Figure 2A). Comparatively, deconvolution of a previously published bulk RNA-seq dataset on the frontal cortex of DOI-or saline-treated mice showed a mix of excitatory and inhibitory cell types, as well as astrocytes (Figure 2B) ^5^. We noted the significant variation of excitatory and inhibitory cell type proportions between samples from each group (Figure 2C). Together, we highlight the selectivity of our RiboTag approach to enrich for single cell types in the mPFC, which allows us to attribute gene expression changes to specific cell types, in stark contrast to bulk RNA-seq.

**Figure 2.**
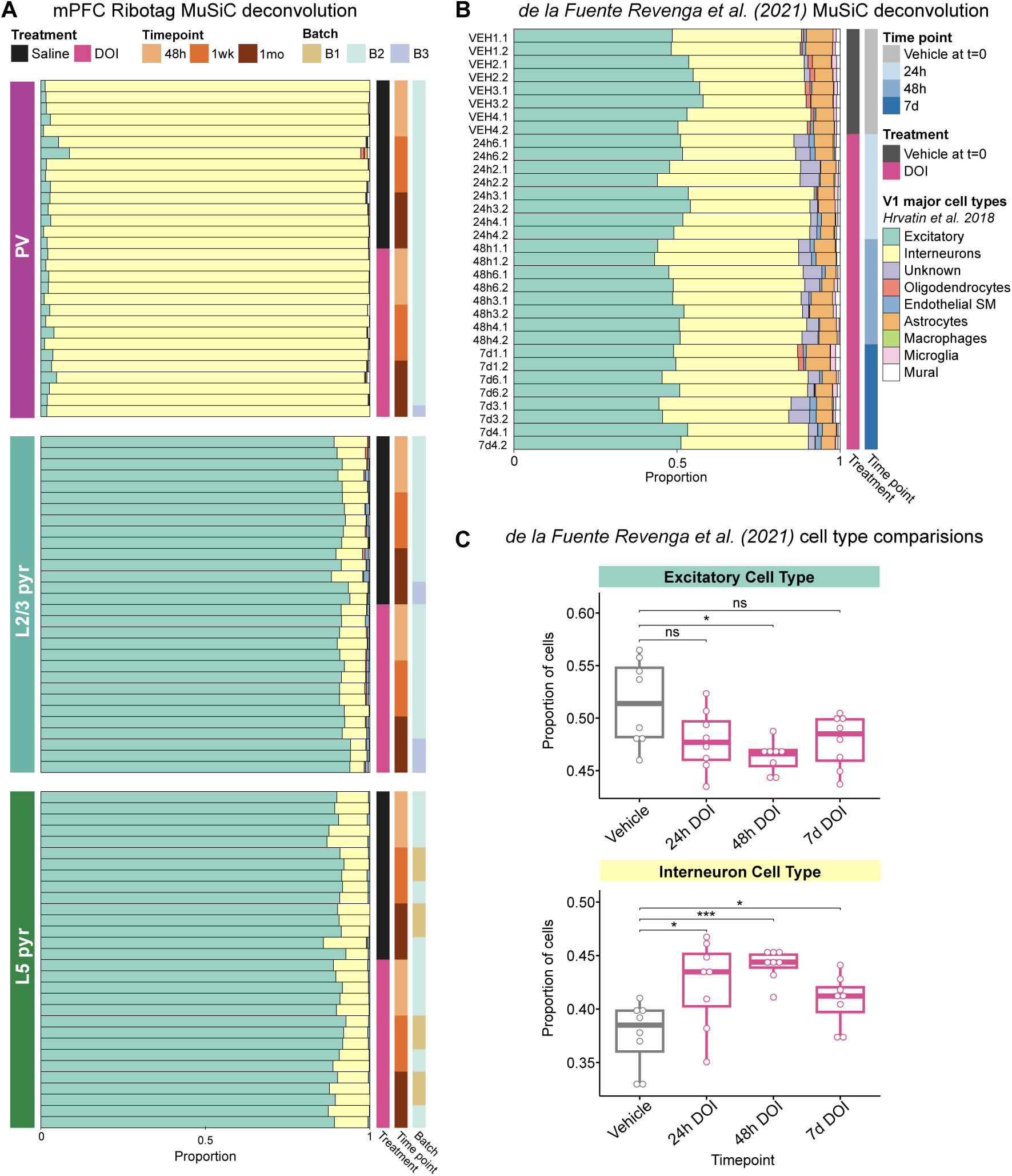
MuSiC deconvolution of bulk RNA-seq datasets. **(A)** Deconvolution results of 90 DOI and saline samples from the current study (30 PV samples, 30 L2/3 samples, and 30 L5 samples) using major cell types from Hrvatin et al., 2018 scRNA-seq dataset (number of cells total = 48,266, number of excitatory cells = 14,287, number of interneurons = 936). **(B)** Deconvolution results of 32 samples in de la Fuente Revenga et al., 2021 RNA-seq dataset (8 samples vehicle, 8 samples 24-hours after DOI, 8 samples 48-hours after DOI, and 8 samples 7-days after DOI). **(C)** Proportion of excitatory and inhibitory cell types represented in de la Fuente Revenga et al., 2021 across treatment conditions. (* *p* < 0.05, *** *p* < 0.001). *p* values are for vehicle versus each DOI time point by unpaired t-test with Benjamini-Hochberg correction.

### Modest changes in gene expression across cell types in the PFC following psychedelic exposure

To explore the longitudinal translatomic changes induced by psychedelics, we performed differential gene expression analysis between psychedelic-treated and saline-matched controls (Figure 3A). Compared to differences between cell types, we found subtle differences in gene expression between treatments. Despite exhaustive analysis using multiple bioinformatic pipelines (EdgeR vs. DESeq2) and cutoffs for statistical significance (q < 0.1 vs 0.05), we observed modest changes in differentially expressed genes (DEGs) across all conditions (Figure S2A and Table S3). Contrary to prior studies that show transcriptional changes occurring in excitatory neurons, we found most DEGs were in PV interneurons, with little to no change in L2/3 and L5 pyramidal neurons, where the magnitude of log_2_FC rarely exceeded 1 (Figures S2B and S2C) ^6,7^. For example, in PV interneurons 48 hours after psilocybin treatment, the immediate early genes (IEGs), Egr1, Egr3, and Fos, are down-regulated (Figure S2B). We noted considerably more DEGs in PV interneurons at one month (Figure S2A). However, these values varied based on the pipeline and exhibited reduced correspondence (Figures S2A and S3A; R^2^ = 0.86 – 1 month DOI vs. R^2^ = 0.94-0.97 other time points). With the log_2_FC values from each pipeline highly correlated to each other across all conditions, we proceeded with values from DESeq2 for the rest of our study (Figures S3A-S3C).

**Figure 3.**
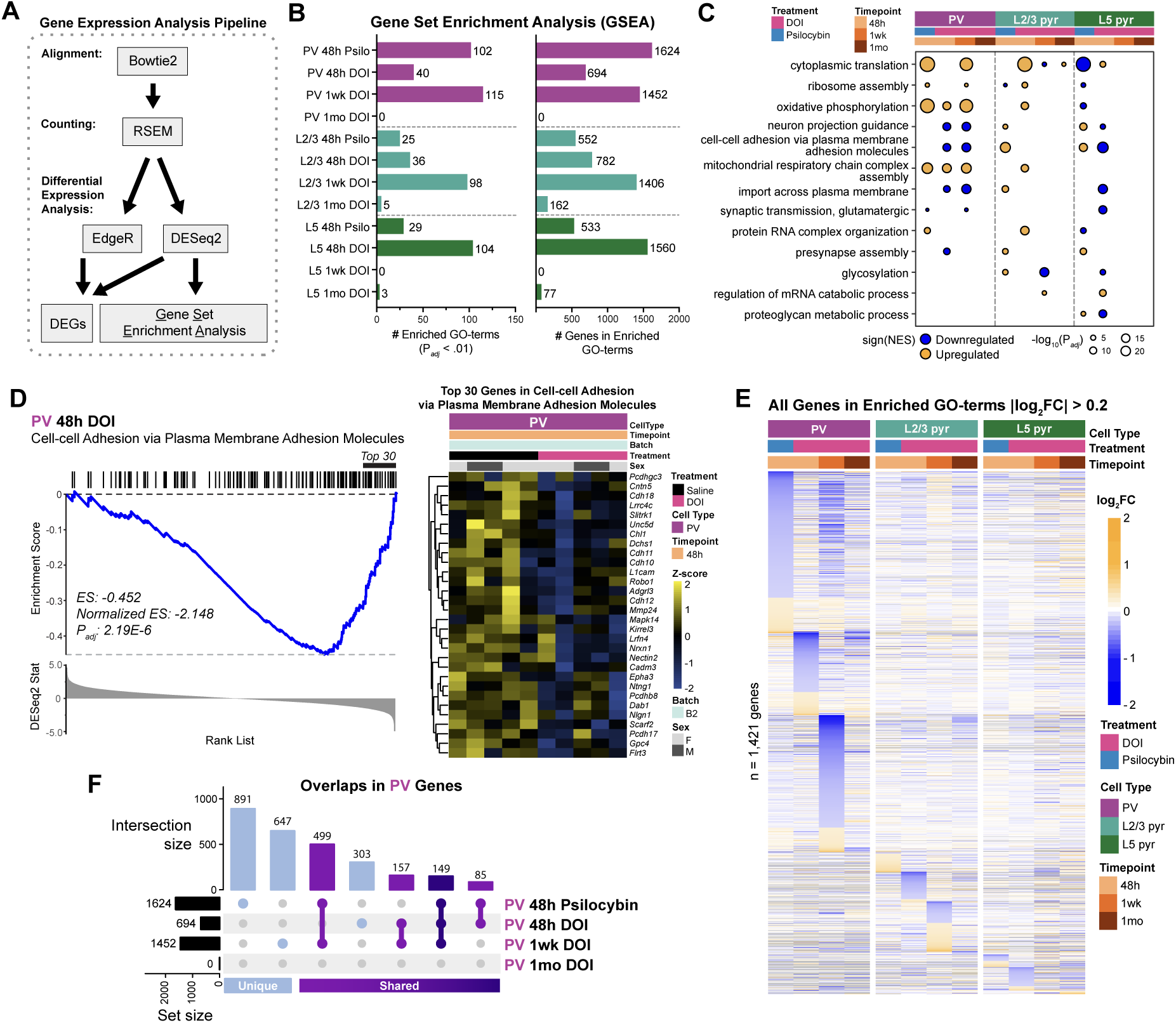
Differential gene expression across cell types in the mPFC following psychedelic exposure. **(A)** Schematic of bioinformatic pipelines used for differential gene expression (DGE) analysis and gene set enrichment analysis (GSEA). **(B)** Number of enriched GO terms from GSEA (p_adj_ < .01), and total number of genes in leading edge of enriched GO terms from each condition. **(C)** Dot plot of top enriched GO terms by p-value, showing one up– and one down-regulated term from each condition by normalized enrichment scores (NES). **(D)** Example GSEA enrichment plot in PV interneurons, 48-hours after DOI, of the GO term, cell-cell adhesion via plasma membrane adhesion molecules. Associated heatmap showing expression of the top 30 genes in the leading edge. **(E)** Heatmap showing fold-change in expression of genes in enriched GO terms (log_2_FC > 0.2 and < –0.2) across conditions. **(F)** UpSet plot showing unique and overlapping genes in enriched GO terms between conditions in PV cells. See also Figures S2, S3, and S4.

Since most genes showed few changes in gene expression, we used Gene Set Enrichment Analysis (GSEA) to examine coordinated changes in groups of genes that share overlapping biological functions (Figure 3A) ^58^. Using GSEA with log_2_FC values from DESeq2 and Gene Ontology (GO) annotations from the Molecular Signatures Database (MSigDB), we identified hundreds of significantly enriched GO terms, with thousands of genes associated with the enriched GO terms (Figures 3B, S4A, and Table S4). With GSEA, we observed that most psychedelic-induced changes occurred at 48 hours and one week, returning to baseline by one month (Figure 3B). Again, changes in PV interneurons exceeded those in both excitatory subtypes regarding the number of enriched GO terms and the log_2_FC magnitude of the genes associated with the GO terms (Figure S4B). There were relatively few overlaps between cell types in the enriched GO terms, suggesting distinct cell type-specific responses to psychedelic treatment (Figure S4C).

Psilocybin or DOI induced cellular responses generalizable to all cell types, such as cytoplasmic translation and ribosome assembly (Figure 3C). Surprisingly, the most significantly enriched GO terms involved metabolic pathways such as oxidative phosphorylation and electron transport chain (Figure 3C and Table S4). We also identified processes important for neuronal functions implicated in psychedelic-induced plasticity, including neuron projection guidance, presynapse assembly, and synaptic transmission (Table S4). For example, in PV interneurons 48 hours after DOI treatment, one of the most significantly down-regulated GO terms was cell-cell adhesion via plasma membrane adhesion molecules, with gene families such as Cadherins, Slits/Robos, and Neurexins/Neuroligns showing coordinated down-regulation (Figures 3D and S4D). Although cell-cell adhesion was also affected in other conditions, they had opposite directions or comprised different genes (Figure 3C and Table S4). In a heatmap showing fold-changes in the expression of genes in enriched GO terms across all conditions, we found that PV interneurons were the most responsive to psychedelics and that these changes were unique to each cell type and time point (Figures 3E, S4B, and S4C). Together, we show that although the changes induced by psychedelics were modest at the gene-by-gene level, with GSEA, we could identify groups of genes that cooperate to induce changes at the pathway level.

### DOI and psilocybin induced distinct molecular pathways at 48 hours

Phenethylamine and tryptamine psychedelics exhibit varying polypharmacology to neuromodulatory receptors examined in our cell type-specific expression analysis (Figure 1E); however, it is unclear whether the two classes of psychedelics induce similar molecular responses in the different cell types. Psychedelics like DOI, LSD, and psilocybin induce the expression of IEGs such as cFOS up to 4 hours after exposure ^11,20,59–61^. To assess whether psychedelics share overlapping patterns at more extended periods as well, we compared the effects of DOI and psilocybin using GSEA of all three cell types 48 hours after exposure. In L2/3 and L5 pyramidal neurons, DOI and psilocybin affected distinct pathways, or in some cases, the same pathway, but in opposite directions (Figure 3C). For example, in L5 pyramidal neurons, psilocybin increased expression of genes associated with GO terms such as synapse formation, neuron projection guidance, cell-cell adhesion, and presynapse assembly. On the contrary, the same GO terms were down-regulated or not affected in DOI at 48 hours (Figure 3C and Table S4). However, in PV interneurons, metabolic pathways were similarly induced by psilocybin and DOI at 48 hours, persisting for up to one week in DOI (Figure 3C). Interestingly, we noted a temporal overlap of genes between psilocybin at 48 hours, and DOI at one week in PV interneurons (Figures 3E and 3F). This shift in the timeline that psilocybin and DOI act on in PV interneurons was not evident in L2/3 or L5 pyramidal neurons (Figures S4E and S4F).

### Robust and persistent alternative splicing changes following psychedelic exposure

Given that we found few changes in gene expression, we hypothesized that regulatory mechanisms beyond transcription may play a role in long-term psychedelic-induced plasticity. We therefore sought to investigate whether alternative splicing, which is distinct but complementary to gene expression and can increase functional diversity ^62,63^, contributes to psychedelic-induced plasticity. Using our high read-depth RiboTag data and rMATs to map splicing junctions, we detected >50,000 alternative splicing events in each cell type and time point. To enrich for differential splicing events more likely to result in functional changes, we filtered out de novo mapped events, events with low reads, and statistically significant events with only less than 10% differences in inclusion levels between treatments (Figure 4A). Using our rigorous criteria, we observed robust changes in alternative splicing compared to gene expression (Figures 4B and S5A). Most differential splicing events were skipped exon (SE) event types (Tables S5-S7). Surprisingly, despite gene expression changes returning to baseline by one month, most alternative splicing changes in PV and L5 occurred one month after DOI exposure. Moreover, the vast majority of genes with at least one differentially spliced event did not overlap with genes identified with GSEA, highlighting distinct molecular programs in gene expression and alternative splicing induced by psychedelics (Figures 4C and S5B-S5D).

**Figure 4.**
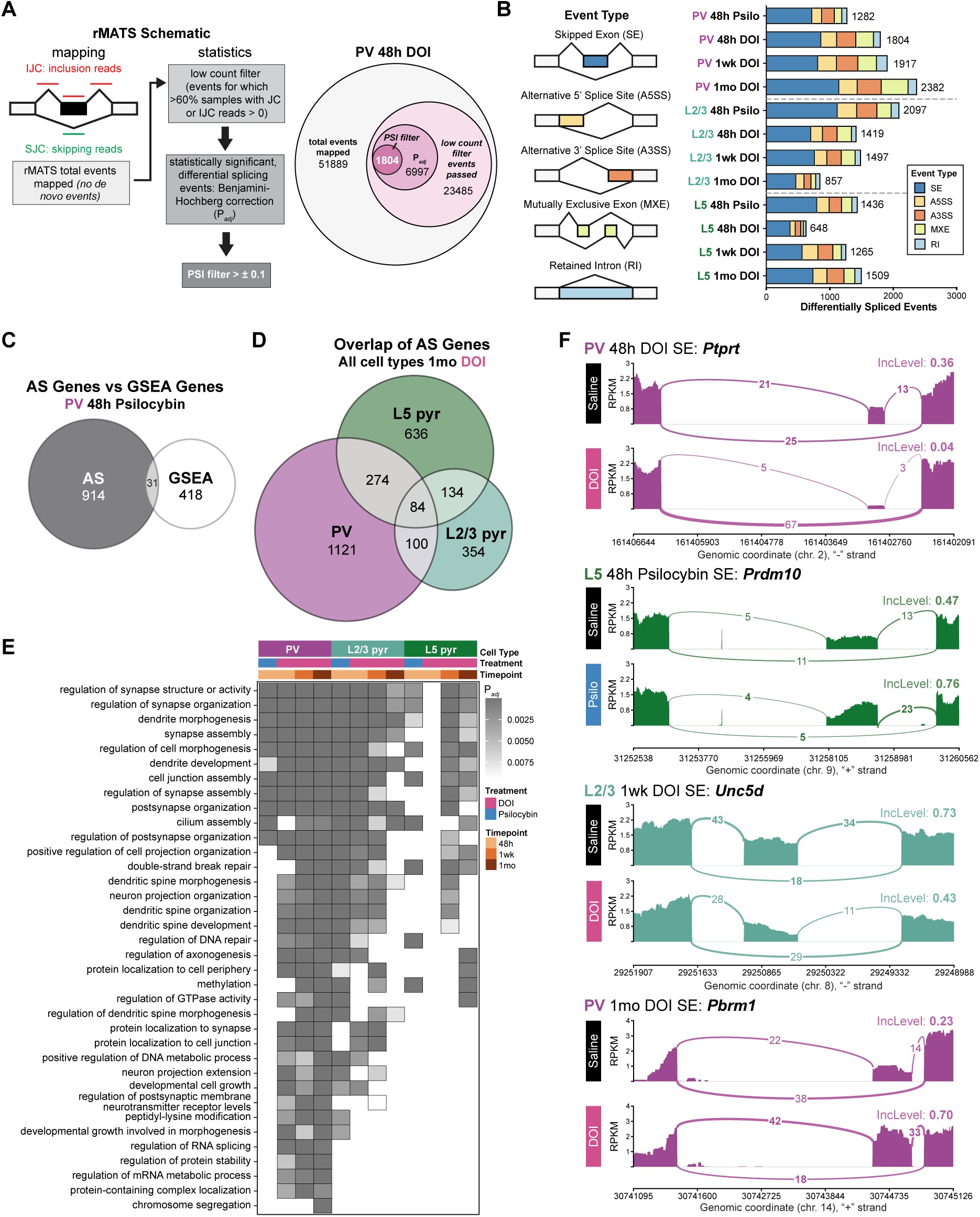
Robust and persistent alternative splicing changes following psychedelic exposure. **(A)** Schematic of rMATs, from mapping of splice junctions to identification of differentially spliced events, with PV interneurons, 48-hours after DOI as an example of the number of events after each filter. **(B)** Schematic of event types (skipped exon, alternative 5′ and 3′ splice sites, mutually exclusive exon, and intron retention), and number of differentially spliced events from all conditions, separated by event type. **(C)** Venn diagram showing overlap of genes identified from alternative splicing analysis vs. GSEA for PV interneurons, 48-hours after psilocybin. **(D)** Venn diagram showing overlap of differentially spliced genes between cell types, 1-month after DOI. **(E)** Tile plot of GO terms for genes with 1 or more differential splicing events, across all conditions. **(F)** Example sashimi plots illustrating read distribution and inclusion level of differentially spliced events from saline-, DOI-, or psilocybin-treated animals. Counts for sashimi plots were generated based on the average read depth and junction-spanning reads within groups. See also Figures S5 and S6.

With psychedelics affecting gene expression in each cell type discriminately, we hypothesized that changes in alternative splicing would also exhibit cell-type specificity. We found cell type-specific expression patterns of bona fide splicing factors in PFC cell types (Figure S6A) ^45^. Coordinately, we identified few overlapping genes that were differentially spliced between cell types across all time points (Figures 4D and S6B). Compared to GSEA, which revealed modest changes to synaptic function at the gene expression level, GO enrichment analysis of differentially spliced genes demonstrated a notable enrichment of synaptic GO terms (total count of synaptic-related pathways identified in all cell types, timepoints, and drugs in GSEA = 52 vs. in AS = 206), with the top GO terms regulating synapse structure or activity, synapse assembly, and dendrite morphogenesis across all cell types (Figure 4E and Table S8). Sashimi plots of differentially spliced genes include *Ptprt*, which is involved in signal transduction, *Prdm10*, and *Pbrm1*, both regulate gene expression, and *Unc5d*, in cell-cell adhesion signaling (Figure 4F) ^64–67^. In contrast to the modest changes in gene expression, we observed large-scale changes in alternative splicing events driving the response to psychedelics, highlighting the importance of examining non-transcriptional processes in understanding the underlying molecular mechanisms of synaptic plasticity.

### DOI reduced abundance of perineuronal nets around L5 but not L2/3 PV interneurons

To link translatomic with functional changes, we examined perineuronal nets (PNNs), specialized extracellular matrix (ECM) structures that regulate cortical plasticity. While PNN accumulation during development coincides with the closure of critical periods and mediates the maturation of cortical circuitry ^68–71^, ECM degradation in the mature cortex can reinstate juvenile-like plasticity ^68,70,72–74^. Recent work shows that psychedelics alter ECM gene expression in the nucleus accumbens during critical period reopening ^9^. However, whether psychedelics change ECM-related structures is unknown. In our mPFC dataset, GSEA identified decreases in extracellular structure organization and cell-matrix adhesion by DOI in PV interneurons by one week (Table S4). In a heatmap showing the fold-change in expression of selected PNN and ECM components across all conditions, we confirmed that the largest change occurs in PV interneurons one week after DOI (Figure 5A).

**Figure 5.**
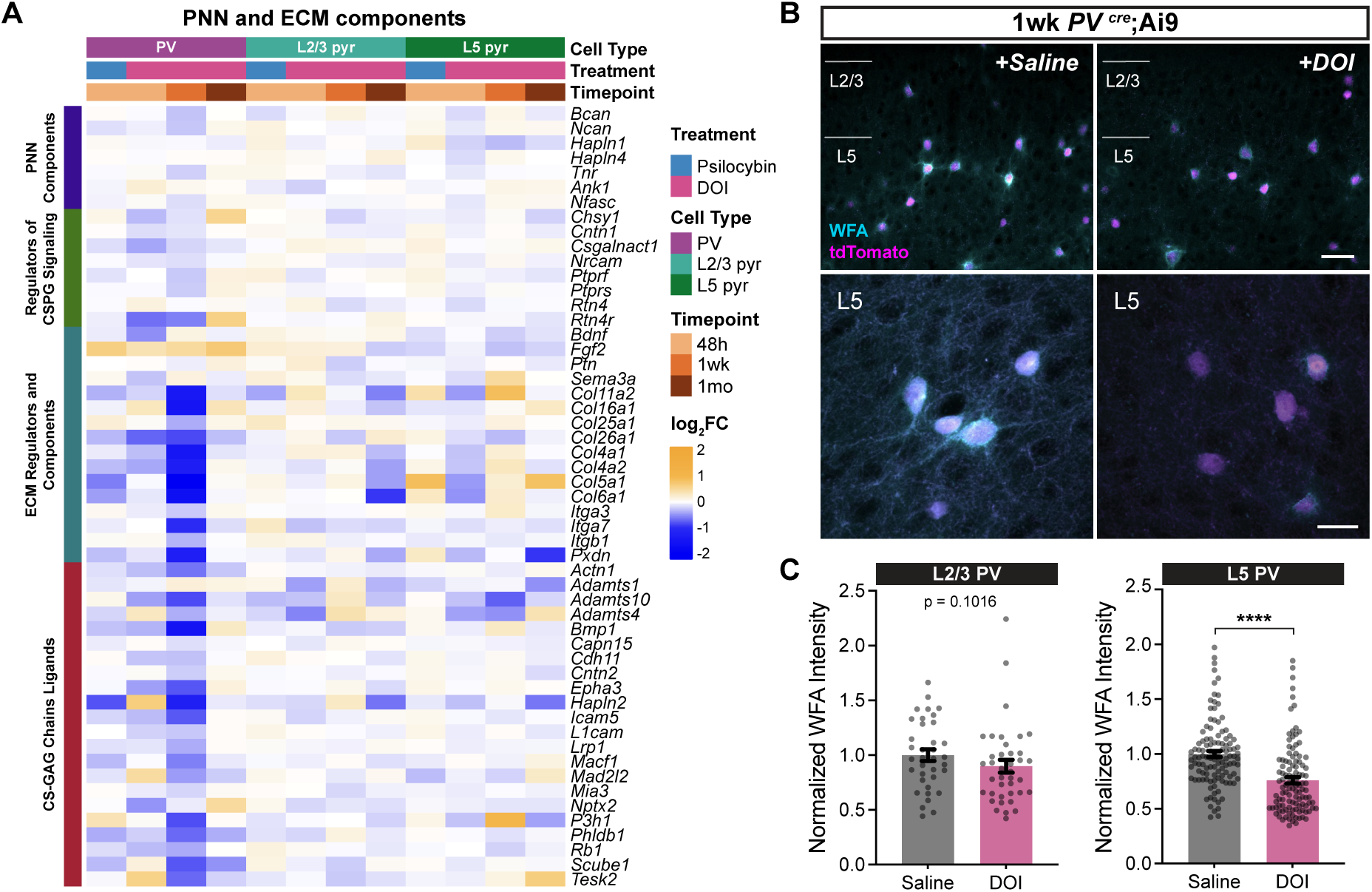
DOI reduces WFA-positive perineuronal nets (PNNs) around L5 PV interneurons. **(A)** Heatmap showing fold-change in expression of PNN and extracellular matrix (ECM) component genes across all conditions. **(B)** Representative images showing WFA labeling surrounding PV interneurons (PV^cre^ x Ai9; tdTomato) in L2/3 and L5 1-week after saline or DOI treatment. Confocal z-projection images of cortical layers at 20x (top) and L5 PV interneurons at 63x (bottom). Scale bars are 50 µm (top) and 20 µm (bottom). **(C)** Normalized WFA intensity surrounding PV interneurons in L2/3 (saline: n=36 cells; DOI: n=39 cells) and L5 (saline: n=112 cells; DOI: n=112 cells). N=3-4 mice per treatment group. (**** *p* < 0.0001). *p* values from permutation test.

To confirm if the downregulation of ECM-related genes corresponds to changes in PNN abundance, we stained psychedelic-treated and saline-matched mPFC with Wisteria floribunda agglutinin (WFA) to label PNNs surrounding PV interneurons ^75–77^. To ensure correspondence with our RNA-seq data, we analyzed neurons expressing tdTomato under the PV-Cre driver. PNNs exhibit distinct localization across cortical layers; thus, we measured WFA intensity separately in L2/3 and L5 of the mPFC ^78,79^. We found a decrease in the average WFA intensity around PV interneurons in L5 but not in L2/3 of the mPFC one week after DOI (Figures 5B and 5C). These results indicate that psychedelics can reduce PNN abundance surrounding PV interneurons in the principal output layer of the mPFC. Importantly, these data confirm that our GSEA reveals biological pathways with functional phenotypes relevant to elucidating the mechanisms of psychedelic-induced plasticity.

### Psilocybin transiently modified the intrinsic and synaptic physiology of PV interneurons

Most translatomic changes occurred in PV interneurons 48 hours after psilocybin treatment (Figure 3B). Since PV interneurons exert substantial control over cortical activity and plasticity ^33,36,78,80–84^, we investigated whether psilocybin altered their physiology. Both GSEA and alternative splicing analysis revealed GO terms that affect synaptic and intrinsic properties of PV interneurons, such as action potential, sodium and potassium ion transport, synapse assembly, and glutamatergic synaptic transmission (Figure 6A). To determine if the psychedelic-induced translatomic changes predict physiological changes, we conducted whole-cell patch-clamp recordings in PV interneurons.

**Figure 6.**
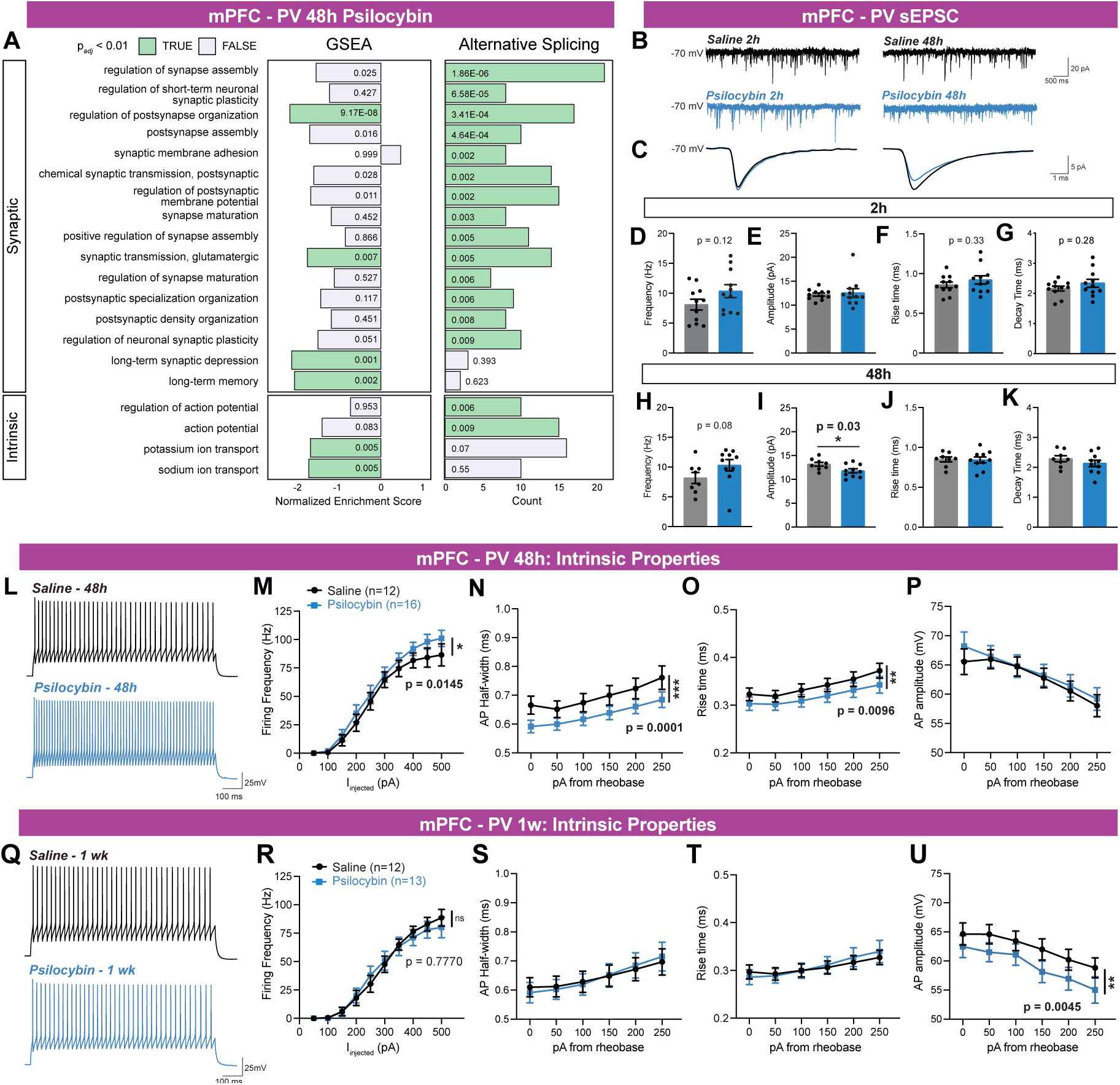
Psilocybin transiently modifies the intrinsic and synaptic physiology of PV interneurons. **(A)** Summary of synaptic and intrinsic physiology related GO terms enriched (padj < 0.01) in alternative splicing analysis vs. GSEA for PV interneurons, 48-hours after psilocybin. **(B-K)** Spontaneous excitatory postsynaptic currents (sEPSC) in saline versus psilocybin treated PV interneurons, 2 hours and 48 hours after treatment. (B) 5s example traces of saline (top) and psilocybin (bottom) at 2-hours and 48-hours. (C) Averaged sEPSC in an example saline and psilocybin treated PV interneuron at 2-hours (left) and 48-hours (right). (D) Frequency, (E) amplitude, (F) rise time, and (G) decay time of sEPSC 2 hours after psilocybin treatment. (H) Frequency, (I) amplitude, (J) rise time, and (K) decay time of sEPSC 48 hours after psilocybin treatment. 2 hours: saline: n = 11 cells; psilocybin: n = 11 cells. 48 hours: saline: n = 8 cells; psilocybin: n = 10 cells. N = 4-6 mice per group. *p* values are for saline versus psilocybin by two-tailed unpaired student’s *t* test or Mann Whitney U test. **(L-P)** Intrinsic properties of PV interneurons 48-hours after drug treatment. (L) Spike trains of example saline (top, black) and psilocybin (bottom, blue) PV interneurons, 48 hours after treatment (200pA injection). (M) Mean firing frequency in response to 1 second somatic current injections. (N) Half-width, (O) rise time, and (P) amplitude of third spike at somatic current injections from rheobase. Saline: n = 12 cells; psilocybin: n = 16 cells. N = 3-4 mice per group. *P* values are for saline versus psilocybin by two-way ANOVA. **(Q-U)** Intrinsic properties of PV interneurons 1-week after drug treatment. (Q) Spike trains of example saline (top, black) and psilocybin (bottom, blue) PV interneurons, 1 week after treatment (200pA injection). (R) Mean firing frequency in response to 1 second somatic current injections. (S) Half-width, (T) rise time, and (U) amplitude of third spike at somatic current injections from rheobase. Saline n = 12 cells; psilocybin n = 13 cells, N = 3 mice per group. *p* values are for saline versus psilocybin by two-way ANOVA. Symbols in bar plots line graphs show mean ± SEM. (* *p* < 0.05, ** *p* < 0.01, **** p < 0.0001). 2h, 2 hours. See also Table 1.

**Table 1.** Electrophysiological properties in PV interneurons after psilocybin treatment. Values reported as mean ± SEM. *p* value from student’s t test or Mann-Whitney U Test. AP kinetics calculated from third spike at 100pA current injection from rheobase. RMP, resting membrane potential. See also Figure 6.

| Property | 48 hours |  |  | 1 week |  |  |
| --- | --- | --- | --- | --- | --- | --- |
|  | *Saline*<br>(n ≥ 9) | *Psilocybin*<br>(n ≥ 14) | *p*<br>*value* | *Saline*<br>(n ≥ 6) | *Psilocybin*<br>(n ≥ 6) | *p*<br>*value* |
| RMP (mV) | -62.22 ± 2.0 | -63.41 ± 2.8 | 0.7521 | -58.5 ± 1.0 | -62.71 ± 1.9 | 0.1577 |
| Input Resistance (MΩ) | 162.6 ± 13.4 | 181.9 ± 18.5 | 0.7988 | 168.0 ± 12.2 | 177.4 ± 24.5 | 0.7457 |
| Membrane τ (ms) | 11.32 ± 0.7 | 15.37 ± 2.9 | 0.3068 | 11.54 ± 1.1 | 17.47 ± 3.2 | 0.1063 |
| Rheobase (pA) | 162.2 ± 21.4 | 138.5 ± 17.1 | 0.3931 | 188.1 ± 19.5 | 174.4 ± 21.5 | 0.6495 |
| AP threshold (mV) | -33.91 ± 1.2 | -33.48 ± 2.4 | 0.8900 | -27.69 ± 1.1 | -29.26 ± 1.3 | 0.3857 |
| AP decay time (ms) | 0.81 ± 0.04 | 0.79 ± 0.05 | 0.8145 | 0.76 ± 0.05 | 0.75 ± 0.04 | 0.9195 |
| fAHP (mV) | -17.66 ± 0.81 | -18.81 ± 1.15 | 0.4472 | -20.91 ± 0.99 | -19.52 ± 0.80 | 0.2865 |
| mAHP (mV) | -18.30 ± 0.82 | -18.80 ± 1.05 | 0.7238 | -21.13 ± 0.99 | -19.76 ± 0.76 | 0.2813 |

Given the enrichment of synaptic GO terms in our AS analysis, including glutamatergic synaptic transmission, we hypothesized that psilocybin at least transiently alters excitatory synaptic transmission onto PV interneurons. We thus assessed spontaneous excitatory postsynaptic currents (sEPSCs) in PV interneurons at multiple time points after psilocybin treatment (Figures 6B-6K). At an acute two-hour time point, psilocybin treatment had no effect on the frequency, amplitude, or kinetics of sEPSCs (Figures 6B-6G). After 48 hours, however, we observed a decrease in sEPSC amplitude in the psilocybin-treated group (Figures 6B, 6C, and 6I). These results indicate that psilocybin induces a reduction in spontaneous glutamatergic transmission in PV interneurons days after exposure.

Moreover, we found that psilocybin treatment induces persistent intrinsic plasticity in PV interneurons. Forty-eight hours after a single dose of psilocybin, we observed a modest increase in firing frequency and a decrease in action potential half-width and rise-time (Figures 6L-6O and Table 1) that returned to baseline by one week (Figures 6Q-6U). The increased intrinsic excitability occurred without significant changes to passive membrane properties or rheobase (Table 1). While there is no change in action potential amplitude at 48 hours (Figure 6P), we observed a decreased amplitude one week after psilocybin treatment (Figure 6U), suggesting that psychedelics alter intrinsic plasticity which varies over time. Together, we show that our translatomic dataset is a valuable resource for assessing functional changes induced by psychedelics.

## Discussion

Previous studies seeking to elucidate the molecular basis of psychedelic-induced plasticity have focused primarily on differential gene expression, overlooking other modes of gene regulation. Here, we present the most extensive study to date (>26 billion reads), assessing long-term and cell type-specific translatomic responses to psychedelic exposure in the mPFC. Our data indicate that psychedelics induce subtle changes in gene expression but robust changes in alternative splicing lasting at least one month after exposure. We establish that different classes of psychedelic drugs, like DOI and psilocybin, engage distinct molecular pathways at differing time scales. We further connect the translatomic changes to their functional correlates by focusing on the relatively small population of GABAergic PV interneurons. Gene ontology terms coincide with our functional changes in extracellular matrix organization and PV interneuron physiology. Together, our findings demonstrate the validity of our dataset as a key resource to understand the cell type-specific effects of psychedelics on the cellular functions in the mPFC.

Unexpectedly, comparing the timeline of our dataset to studies that find month-long changes ^32^ in dendritic spines in L5 pyramidal neurons in the mPFC, we observed modest changes in gene expression across three cell types in the mPFC from 48 hours to one month after DOI or psilocybin. The few changes in gene expression that we observed were of low magnitude (<1 log_2_FC) across all genes and time points. These findings are inconsistent with previously published bulk RNA-seq data in the PFC, which found substantial changes in gene expression at 24 hours, 48 hours, and one week after DOI treatment ^5^. Other cell type-specific studies using snRNA-seq find significant changes in excitatory neurons from one hour to 72 hours after psilocybin exposure, with few or no changes in interneurons such as PV and SST ^6,7^. In contrast, most changes in gene expression that we identified were in PV interneurons, with no change in L5 pyramidal neurons. We attribute the inconsistency in findings to the experimental designs in the respective studies, including differences in the use of saline-matched controls and sequencing techniques ^5–7,10^. Our deconvolution analysis suggests that previously reported bulk RNA-seq studies have an unbalanced proportion of inhibitory and excitatory cell types across samples, especially between treatments. Comparatively, our RiboTag approach allows us to examine a single genetically-defined cell type, focusing on the translatome instead of the transcriptome, in which we retain mRNAs in axons and dendrites that are lost during nuclei isolation ^53,54^. Moreover, inhibitory cell types like PV interneurons represent smaller clusters and fewer neurons than excitatory neurons, presenting challenges in identifying modest gene expression changes using snRNA-seq or scRNA-seq ^85^. Moreover, other studies report that transcriptomic changes peak at one hour and return to baseline by 24 hours based on GO term analysis ^7^. The more modest changes in gene expression that we observed may be due to our assessment of expression at later time points, days to a month after psychedelic exposure.

Despite differences in the magnitude of gene expression, we identified changes in gene expression previously implicated in psychedelic-induced plasticity. Expression of IEGs, such as cFOS and Egr1, decreases 48 hours after psilocybin, suggesting homeostatic regulation days after psychedelic exposure ^5,11,20^. Using GSEA to examine coordinated changes in gene expression, we identified biological pathways affected by psychedelics, including synaptic transmission and axon growth, which recapitulate prior functional studies. We further expand on temporal features of long-term psychedelic action and report that changes in gene expression return to baseline after a month.

Although classic serotonergic psychedelics are agonists of the 5HT_2A_R, we find that psychedelics from the tryptamine and phenethylamine classes induce distinct changes at the molecular level 48 hours post-psychedelic exposure in all three cell types^11,19–24^. Interestingly, in PV interneurons, we observe an overlap between psilocybin after 48 hours and DOI after one week, suggesting that psilocybin and DOI may engage overlapping molecular pathways that are shifted temporally. These findings raise the possibility that the different molecular responses observed between classes of psychedelics at single time points may reflect their different timescales of action rather than fundamentally distinct effects. Previous research suggests that although psychedelic compounds operate on distinct timescales, their capacity to induce plasticity is a common feature^9^. Understanding the functional impact of psychedelics, therefore, requires careful consideration of the timescale of action for each compound. These findings emphasize the importance of avoiding considering psychedelics as a single entity.

Critically, while existing transcriptomic studies of psychedelic-induced plasticity predominantly focus on differential gene expression, we used RiboTag deep sequencing to obtain deep coverage of splice junctions, enabling the analysis of alternating splicing. Not only are splicing changes more abundant and synapse-specific compared to gene transcription, they last up to one month after a single dose of psychedelics, coinciding with structural changes observed at dendrites ^32^. Importantly, genes with differential splicing are distinct from those with differential gene expression. These results suggest that psychedelics robustly engage mechanisms that change alternative splicing and contribute to long-term phenotypic changes after psychedelic exposure. Together, our GSEA and alternative splicing analysis provide a comprehensive picture of the persistent gene regulatory responses to psychedelic exposure.

To validate whether our translatomic analysis identifies biological pathways that are functionally altered by psychedelics, we focused our efforts on PV interneurons. Recent evidence indicates that psychedelics acutely activate mPFC PV interneurons and that the activation of 5HT_2A_R-expressing PV interneurons in other brain regions promotes the acute anxiolytic effects of psychedelics ^8,14^. While inhibition is disrupted in the mPFC in mood disorders and altering inhibitory activity has been suggested as a mechanism of action for fast-acting antidepressants, previous studies primarily focus on psychedelic modulation of excitatory transmission in principal neurons ^5,9,32,86–91^. Our study shows that more changes in gene expression are found in PV interneurons, compared to principal pyramidal neurons in L2/3 and L5 of the mPFC. Interestingly, our analysis of serotonin receptor distribution in the mPFC indicates that PV interneurons express higher levels of 5HT_2A_R than L5 pyramidal neurons, a cell type previously reported to be enriched with these receptors ^26^. Our findings indicate that PV interneurons are an understudied yet important component of the psychedelic response.

In PV interneurons, GSEA identified several biological processes affected by DOI and psilocybin, including metabolism, synaptic transmission, ion transport, and extracellular matrix structure. While individual ECM-related genes did not meet the threshold to be considered DEGs, GSEA revealed collective alterations in ECM gene expression that corresponded to a reduction in PNN abundance surrounding L5 PV interneurons one week after treatment with DOI. While previous studies suggest a role for ECM remodeling in the psychedelic response, we provide the first evidence that psychedelics can induce remodeling of the ECM, evident long after a single exposure ^9^. Considering that PNN reduction enhances cortical plasticity and reopens critical periods, our findings offer further support for the hypothesis that psychedelics induce a prolonged and heightened phase of plasticity that persists well beyond the presence of the drug ^9,68,73,92^. Our GSEA was thus critical to reveal biological pathways positioned to play a role in long-lasting psychedelic plasticity, which may be undetected using standard DEG analysis.

While our gene expression analysis highlights functional effects of psychedelics, our alternative splicing analysis revealed a broader range of gene ontology terms impacted by psychedelic exposure. Notably, we found an enrichment of synaptic-related terms in our differentially spliced genes. In PV interneurons, 48 hours after psilocybin treatment, AS and GSEA analyses identified changes to mRNA involved in synaptic physiology, action potential regulation, and ion transport. Concurrent with these RNA changes, we observed a decrease in the magnitude of spontaneous excitatory postsynaptic currents, an increased firing rate, and altered action potential dynamics. Interestingly, the changes in synaptic excitation we observed 48 hours after psilocybin treatment were not evident at two hours, suggesting that these physiological changes were not an immediate response to psilocybin. Similarly, given that the effects of psychedelics on intrinsic physiology differ in the days to a week after exposure, it is likely that long-term psychedelic-induced plasticity relies on gene regulation that evolves over time. Although both GSEA and AS analysis identified functional effects of psychedelics in neuronal physiology, we found that the majority of the changes in synaptic transmission-related mRNAs were associated with alternative splicing. Our findings support previous studies reporting PV interneuron plasticity without significant gene expression changes and highlight the need to examine non-transcriptional gene regulation in cortical plasticity ^93^.

The regulatory mechanisms that enable neural plasticity for learning while stably retaining existing memories throughout a lifetime remain unclear. Our findings demonstrate that a single psychedelic dose robustly and persistently alters alternative splicing lasting a month with modest changes in gene expression. These results suggest that alternative splicing can expand molecular complexity for cell-to-cell signaling without disrupting gene expression patterns needed for neuronal homeostasis. Pyramidal neurons and PV interneurons exhibit similar magnitudes and durations of psychedelic-induced alternative splicing changes. However, each cell type expresses unique differentially spliced genes, demonstrating that alternative splicing is widely utilized and engages distinct cell type-specific plasticity responses. It is unclear which mechanisms lead to long-term changes in alternative splicing, though histone modifications that flank alternative exons are attractive candidates ^47,94^. Future efforts to ascertain how psychedelic-induced alternative splicing events impact neural circuitry and behavior will be critical. In conclusion, our study highlights that understanding the molecular elements contributing to long-term psychedelic effects requires insights into mechanisms beyond gene transcription and provides an appealing hypothesis for how neural function remains stable in the face of enduring change.

## Limitations of the Study

Although our study includes both male and female mice, a larger sample size is needed to statistically compare sex-dependent psychedelic responses. Nonetheless, our dataset is the most extensive, high-read depth RNA-sequencing study of the temporal response of isolated cortical neuron types to psychedelic treatment. With our rich dataset, we foresee the discovery of other psychedelic-related targets and pathways, and although we were constrained by the article length, we highlighted the effects of psychedelics on PNNs and PV physiology. We expect future studies to utilize this dataset to explore novel mechanisms engaged by psychedelics in contributing to their therapeutic action.

## Resource Availability

### Lead contact

Further information and requests for resources should be directed to Dr. Andrea Gomez.

### Materials available

This study did not generate new unique reagents or materials.

### Data and code availability

Data and code are on embargo and will be made publicly available following peer review.

## Acknowledgements

We thank S.J. Burden, M. Feller, H. Adesnik, and N. Ingolia for insightful comments on the manuscript and expert advice from E. Furlanis. Y.H. was financially supported by a National Institutes of Health T32 grant (5T32GM007232) and a Predoctoral Fellowship from the PhRMA Foundation. A.S.T. was supported by a National Institutes of Health T32 grant (5T32NS095939). E.V. was supported by a Marco Antonio Firebaugh Scholarship. This work was supported by funds to A.M.G. by a NARSAD Young Investigator Grant from the Brain & Behavior Research Foundation, the Shurl and Kay Curci Foundation, the Rennie Fund for the Study of Epilepsy, a Rose Hills Innovator Award, a IDOR-Innovative Genomics Institute grant, an Alfred P. Sloan Research Fellowship, a McKnight Scholar Award, and from anonymous family donations to the UC Berkeley Center for the Science of Psychedelics. Confocal imaging experiments were conducted at the CRL Molecular Imaging Center (RRID:SCR_017852) supported by the Helen Wills Neuroscience Institute.

## Author contributions

Y.H., M.A.S.F., A.S.T., A.M.G. conceived and designed the experiments. Y.H. conducted the RiboTag experiments. M.A.S.F. and Y.H. performed the RNA-seq data analysis. A.S.T. conducted and analyzed the histology and physiology experiments. E.V. performed histology experiments. Y.H., M.A.S.F., A.S.T., and A.M.G. wrote the manuscript. All authors reviewed the manuscript before submission.

## Methods

### Drug

Psilocybin was obtained from the National Institute of Drug Abuse (NIDA) drug supply program. DOI (Sigma-Aldrich, D101) and psilocybin were dissolved in 0.9% saline and administered at a dose of 2mg/kg intraperitoneally (i.p).

### Mice

Rpl22-HA (RiboTag) mice, Ai9 tdTomato mice, and PV-Cre mice were obtained from Jackson Laboratories (Jax stock no: 011029, 007909, 008069, respectively). Rbp4-Cre mice and Cux2-CreERT2 mice were obtained from the Mutant Mouse Resource and Research Center (MMRRC stock no: MMRRC_036400-UCD, MMRRC_032779-MU, respectively). All lines were maintained on a C57Bl6/J background. Animals were group housed in a reversed 12-h light/dark cycle, with food and water ad libitum. Experiments were performed when animals were 8-15 weeks old and randomly assigned to experimental groups. All experiments were performed during the dark phase. All experiments were performed in accordance with the regulations of the Institutional Animal Care and Use Committee (IACUC) of the University of California, Berkeley.

### Method Details

#### Tamoxifen induction

Cux2-Cre^ERT^ mice were induced with a single dose of tamoxifen (0.2mg/g body weight, 20 mg/mL solution, Sigma Aldrich, T5648) dissolved in corn oil, and administered by oral gavage (PO) at postnatal day P14.

#### Immunohistochemistry and WFA staining

Animals from postnatal weeks 8-11 were transcardially perfused with 4% paraformaldehyde in PBS. The brains were fixed overnight at 4°C and washed three times in PBS. 50μm coronal slices were cut using a vibratome (Leica Microsystems VT1000 S). Slices were blocked for 1 hour in .05% TritonX-100 and 1% BSA blocking solution. For RiboTag validation experiments, slices were incubated with primary antibody (rat anti-HA monoclonal clone 3F10 (Roche, 12158167001; 1:500)) in blocking solution overnight at 4°C. Slices were then washed five times in PBS containing 0.05% Triton X-100, followed by incubation for 1 hour at room temperature with secondary antibody (Streptavidin, Alexa Fluor™ 488 Conjugate (Molecular Probes, S32354; 1:500)) in blocking solution, and washed five times in PBS before mounting onto microscope slides with DAPI Fluoromount-G (SouthernBiotech, 0100–20). For perineuronal net experiments, slices were incubated for 2 hours at room temperature with Wisteria Floribunda Lectin Fluorescein (WFA; Vector Laboratories, FL-1351; 1:1000)) and Hoescht (Bio-Techne, 33342; 1:1000) in blocking solution and washed five times in PBS before mounting onto microscope slides with Fluoromount-G (SouthernBiotech, 0100–01).

#### RNA isolation by RiboTRAP pulldowns

RNA purification from RiboTag mice were modified from previously published protocols ^95^. Brains from male and female mice between postnatal weeks 8-10 were removed and stored in ice-cold PBS. To dissect the PFC, a brain was first placed into an acrylic 1mm brain matrix (Stoelting 51380). The brain was sliced coronally with razor blades from Bregma 3 to Bregma –1. The resulting section was cut again in a fan-shape above the corpus callosum, and medial to the motor cortex, in order to isolate the medial PFC. Cortices were then lysed in 1mL of supplemented homogenization buffer (50mM Tris-HCl pH 7.4, 100mM KCl, 12mM MgCl2, 1mg/mL heparin (Sigma Aldrich, H3393), 1x cOmplete, mini, EDTA-free protease inhibitor cocktail (Roche, 04693159001), 200 units per ml RNasin Ribonuclease inhibitor (Promega, N2115), 100 μg/mL cycloheximide (Fisher Scientific, 357420010), and 1 mM dithiothreitol (DTT; Fisher Scientific, 327190010) in DEPC-treated water). Cortices were homogenized in a 7mL Dounce tissue grinder (Sigma Aldrich, D9063). The lysate was centrifuged at 2,000 x g for 10mins, and the supernatant transferred to a new tube. IGEPAL CA-630 (Sigma Aldrich, I8896) was added to a final concentration of 1% and incubated on ice for 5mins. The lysate was centrifuged at 13,000 x g for 10mins, and the supernatant transferred to a new tube. For whole cell lysate (WCL), 10% volume of the lysate was taken and resuspended in 350uL RLT buffer from RNeasy Mini kit (Qiagen 74106) with 2-mercaptoethanol (Fisher Scientific, O3446I-100). For immunoprecipitation (IP), 50uL of anti-HA beads (Fisher Scientific, 88837) pre-washed with supplemented homogenization buffer were added to the remaining supernatant and incubated at 4°C for 3-4 hours with rotation. After incubation, beads were washed 3 times in high salt wash buffer (50mM Tris-HCl pH 7.4, 300 mM KCl, 12mM MgCl2, 1% IGEPAL CA630, 100 μg/mL cycloheximide, and 1 mM DTT in DEPC-treated water). Beads were then eluted in 350 μL of RLT buffer with 2-mercaptoethanol. RNA purification was performed using a RNeasy Mini kit following the manufacturer’s instructions.

#### Reverse transcription and RT-qPCR

20ng of RNA for PV samples and 50ng of RNA for Rbp4 and Cux2ERT samples were used for reverse transcription with SuperScript III First Strand Synthesis SuperMix (Invitrogen, 11752250). RT-qPCR was performed with Power SYBR Green PCR Master Mix (Applied Biosystems, 4368706) on a QuantStudio 3.

#### Library preparation and Illumina sequencing

Purified RNA samples were submitted to Novogene Corporation for sequencing. Sample QC was performed and only samples with RNA integrity number (RIN) values >7 were used for subsequent steps. 50 ng of RNA was used for library preparation with TruSeq stranded RNA library prep kit with poly-A enrichment. The libraries were sequenced 150bp paired-end on a NovaSeq 6000 platform for DOI samples, and NovaSeq X Plus for psilocybin samples, at 200 million reads per sample.

#### Electrophysiology

Two hours, 48 hours, and one week after intraperitoneal injection, PV-Cre;Ai9 mice aged 10-15 postnatal weeks were anesthetized with isoflurane and then transcardially perfused with chilled dissection buffer (in mM): 75 sucrose, 87 NaCl, 2.5 KCl, 1.25 NaH_2_PO_4_, 0.5 CaCl_2_, 7 MgCl_2_, 25 NaHCO_3_, 1.2 ascorbic acid, and 10 dextrose, pH 7.3-7.4. Coronal prefrontal cortex slices (300μm) were prepared with a vibratome (Leica VT1200 S) in chilled oxygenated (95% O_2_/5% CO_2_) dissection buffer. Slices were transferred to oxygenated dissection buffer at 32°C for approximately 30 min and then transferred to artificial cerebrospinal fluid (ACSF; in mM): 124 NaCl, 2.5 KCl, 1.25 NaH_2_PO_4_, 2.5 CaCl2, 2 MgSO_4_, 26 NaHCO_3_, 10 dextrose. Slices incubated at room temperature at least 30 min to allow for recovery, then transferred to the recording chamber and perfused (1.5-2.5 ml/min) with oxygenated ACSF at room temperature. Somatic whole-cell recordings were performed in mPFC. tdTomato+ parvalbumin interneurons were selected by fluorescence illumination (CoolLED). Cells were current or voltage clamped with Multiclamp 700B amplifier (Molecular Devices) and digitized by Digidata 1550B (Molecular Devices). Patch pipettes (4-8 MΩ) were filled with current clamp (I_c_) solution or voltage clamp (V_c_) solution (in mM: I_c_ = 135 K-gluconate, 5 NaCl, 5 Mg-ATP, 0.3 Na-GTP, 10 Na-phosphocreatine, 10 HEPES, 100 Alexa Fluor 488; V_c_ = 135 CsMeSO_3_, 10 HEPES, 8 NaCl, 4 Mg-ATP, 0.3 Na-GTP, 0.3 EGTA, 2 Qx314-Cl, 0.1 spermine). Data were digitized at 20 kHz with Clampex 11 (Molecular Devices).

To measure intrinsic properties, cells were current-clamped to hold membrane potential at –70mV and data were analyzed using Clampfit 11, Easy Electrophysiology (v2.7.3), and Python pyABF. Resting membrane potential and input resistance were calculated using pyABF code adapted from Spikes and Bursts blog ^96^ and reported as the average of 2-3 technical replicates. Firing rate and action potential kinetics were calculated with Easy Electrophysiology software (dV/dT = 20mV/ms) and reported as the average from 2-3 technical replicates. Action potential kinetics were measured from the third spike. Membrane tau was calculated in Easy Electrophysiology and reported as an average of 20 traces per cell. To measure sEPSC, two minutes of voltage clamp were recorded at –70 mV, respectively. Holding potentials were adjusted for liquid junction potential (+7mV). Series resistance was not compensated. Postsynaptic current data was low-pass Bessel filtered at 2kHz. Amplitude, frequency, and kinetics of sPSCs were quantified by template-based analysis in Easy Electrophysiology software with an amplitude threshold of 5pA. For all experiments, cells were excluded that required current injection with a magnitude greater than 300pA to hold at –70mV or that had an access resistance > 25 MΩ during the experiment. All recordings and analysis were performed blind to treatment.

### Quantification and Statistical Analysis

#### RNA-seq data processing and QC

Data quality assessment and sequence read primer trimming were performed by FastQC v0.11.9 and Cutadapt 3.5, respectively. We considered trim_galore version 0.6.7 with *--phred3*3 parameter. The data was then aligned to a mouse reference genome build (Mus musculus; GRCm39: RefSeq assembly accession = GCF_000001635.27) using RSEM v1.3.1 ^97^. Bowtie2 version 2.4.4 aligner index was created using *rsem-prepare-reference –-bowtie2* command and a gene-level read count matrix was generated using *rsem-calculate-expression –-bowtie2* command. Gene-level counts of each sample were combined using *rsem-generate-data-matrix* command.

#### Single-cell deconvolution analyses

To compare the profile of the single-cell brain atlas and the bulk-cell population specimen of this and de la Fuente Revenga studies, we used the MuSiC version 1.0.0 method ^57^. We downloaded datasets from GEO: GSE102827: mouse visual cortex single-cell atlas ^56^; GSE161626: RNA-seq on neurons (NeuN+) of the mouse cortex frontal lobe ^5^. We considered the eight major class cell type annotations, totaling 48,266 cells across nine samples to train the model. The gene signature was based on the ∼14,000 common genes expressed in both scRNA-seq and bulk RNA datasets.

#### Differential gene expression analyses

Differential gene expression and functional enrichment analysis were performed using R version 4.4.1, Bioconductor version 3.19, and packages therein. EdgeR version 4.2.2 was used to normalize and perform differential expression analysis on the RSEM quantified reads. DESeq2 version 1.44.0 was used to normalize, perform differential expression analysis on the RSEM quantified reads, and prepare the data for functional enrichment analysis using fgsea 1.30.0 ^98^. The sorted stat values rank were considered as *stats* parameters along with *minSize = 15* and *maxSize = 200*. We considered Gene Ontology (GO) annotations from the Molecular Signatures Database (MSigDB) and subset the candidate pathways to the genes that are present in our data to avoid bias. For both EdgeR and DESeq2 pipelines we filtered genes considering minimum count equals 50 and only protein-coding genes annotations from biomaRt 2.60.1.

#### Alternative splicing analysis

For alternative splicing analysis, we first applied the same procedures previously described for assessing data quality and sequence read primer trimming. The reads were then aligned to a mouse reference genome build (Mus musculus; GRCm39: RefSeq assembly accession = GCF_000001635.27) using STAR 2.7.10a with parameters *--outFilterMatchNmin 50 –-outFilterMismatchNmax 100*. Differential alternative splicing (AS) events were obtained using rMATS turbo v4.3.0 and Python 3.10.9 with parameters *--readLength 150 –-variable-read-length –-allow-clipping* ^99^. De novo events were not considered. We filtered out events with zero counts on 60% or more of the samples in each treatment group and calculated adjusted p-value for the remaining events considering Benjamini & Hochberg (BH) method. Analyses described in the main results used only events with rMATS P-value adjusted < 0.05, and |ΔΨ| > 0.1 in treatment versus control RNA-seq sample groups.

#### Batch correction for splicing analysis

To adjust for batch count effects across skipping and inclusion junction reads we used an empirical Bayes framework (comBat), available in sva version 3.52.0 ^100^. First, we obtained the inclusion junction and skipping junction raw counts by running rMATs with *--statoff* parameter. We filter out “noise” events if their values were zero on 80% or more of the samples. The remaining events were corrected using comBat and the transformed count values were rounded to integer. Negative values were flatted to zero. Next, final rMATS count files were formed using the corrected junction counts and the “noise” junction counts events. These files were then informed back to run rMATs statistical model with *--task stat* parameter. Additionally, we performed the procedures described in the alternative splicing analysis session without considering batch correction and filtered out batch-corrected events that were overlapping non-corrected events.

#### Perineuronal net intensity analysis

Perineuronal nets (PNNs) were quantified by measuring WFA fluorescence surrounding individual tdTomato^+^ cells in layer 2/3 and 5 of the mPFC of PV^cre^;Ai9 mice. 20x objective images were acquired in a single imaging session using laser scanning confocal microscopy (LSM880, Zeiss) with consistent laser power, gain, brightness, contrast, and z-plane interval. For each animal, cells were analyzed at the following Bregma positions: +0.73, and +0.25 ^101^. Cortical layers were defined as follows: 100-225µm from pial surface for L2/3 and 250-450µm from pial surface for L5. The experimenter was blind to treatment and sex, and randomly selected tdTomato^+^ cells for analysis without viewing WFA staining. The z-plane center of each cell was identified and the integrated density of WFA was summed in an ROI (25×25µm) ± 6 z-planes from the center of each cell (0.50µm z-plane interval). ROIs that overlapped with other tdTomato^+^ cells in this z-plane range were excluded. Cells were randomly selected from each tissue slice to balance sample size by animal, bregma, and treatment, and then pooled by treatment group for statistical analysis. WFA intensity was normalized to the mean of the saline group in each layer. Normalized means between treatment groups were compared by permutation test with α = 0.05 (Python).

#### Electrophysiology

Statistical analysis was performed using Graph Pad Prism (v.10.3.1). Two-way ANOVA was used to compare means for x-y data. To compare individual measurements, data were tested for normality using the Shapiro-Wilk test. For normally distributed data, two-tailed unpaired Student’s *t* tests were used to compare means. The Mann-Whitney U test was used for non-normal data.

## Supplemental Information

Document S1. Figures S1-S6.

Table S1. Metadata and QC analysis of sequencing data for all RiboTag samples, related to STAR Methods

Table S2. Differentially expressed genes between cell types from all saline samples using DESeq2, related to Figure 1

Table S3. Differentially expressed genes between treatments across all conditions using DESeq2, related to Figure 3

Table S4. Enriched GO terms from gene enrichment set analysis (GSEA), related to Figure 3

Table S5. rMATS output of alternative splicing events associated to PV, related to Figure 4

Table S6. rMATS output of alternative splicing events associated to L2/3, related to Figure 4

Table S7. rMATS output of alternative splicing events associated to L5, related to Figure 4

Table S8. GO terms associated with differentially spliced genes across all conditions, related to Figure 4

---

**Figure S1.**
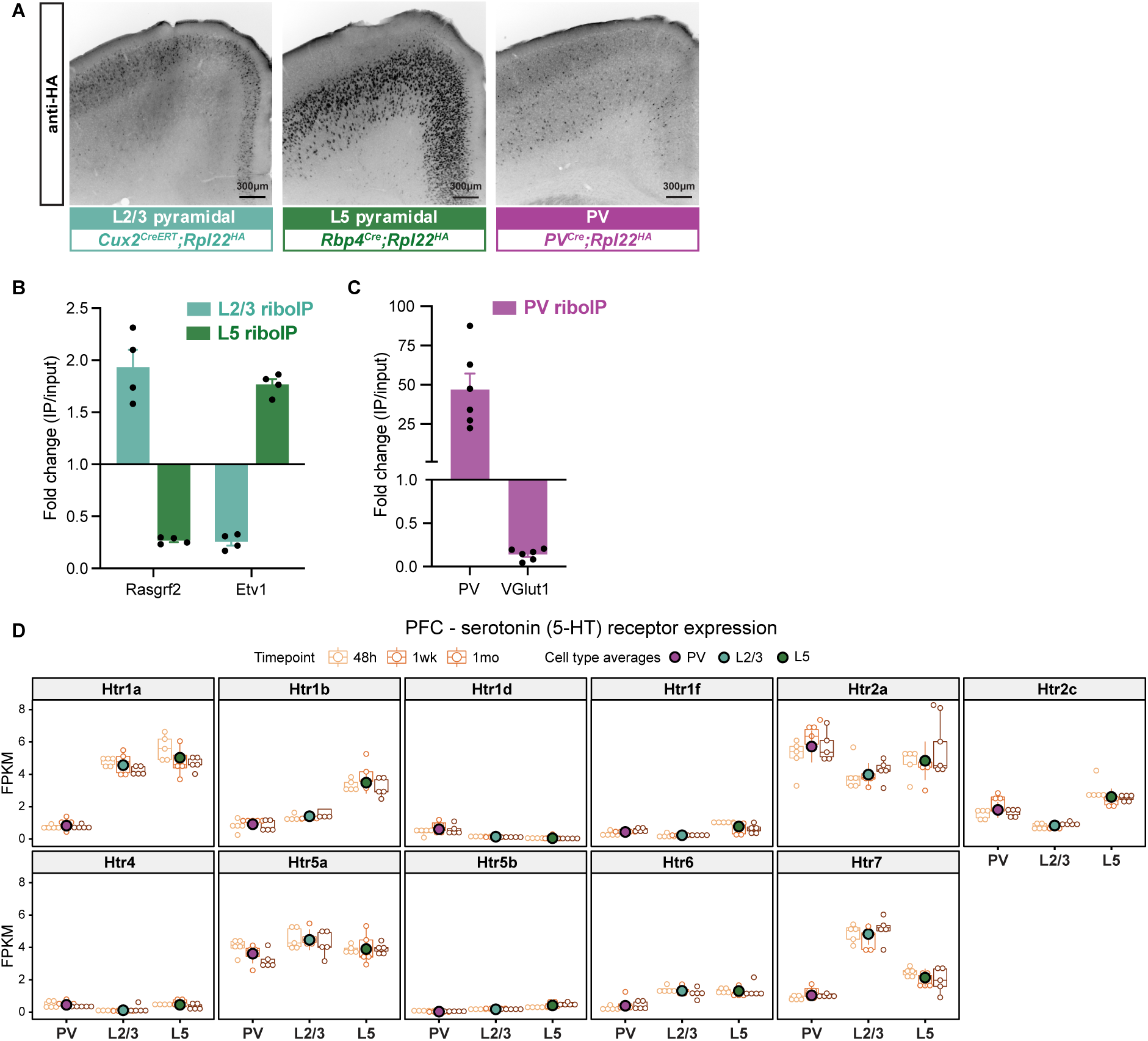
Expression of cell markers and serotonin (5-HT) receptors in RiboTag cell types from the mPFC, related to Figure 1. **(A)** IHC staining of HA-tagged ribosomes in the 3 Cre drivers (Cux2-Cre^ERT^ for L2/3 pyramidal neurons, Rbp4-Cre for L5 pyramidal neurons, and PV-Cre for PV interneurons). **(B)** RT-qPCR of purified RNAs from Cux2-Cre^ERT^ and Rbp4-Cre against L2/3 and L5 cell type-specific markers Rasgrf2 and Etv1, respectively. **(C)** RT-qPCR of purified RNAs from PV-Cre against inhibitory and excitatory markers PV and VGlut1, respectively. **(D)** FPKM of 5-HT receptors across cell types in all saline samples, separated by time point.

**Figure S2.**
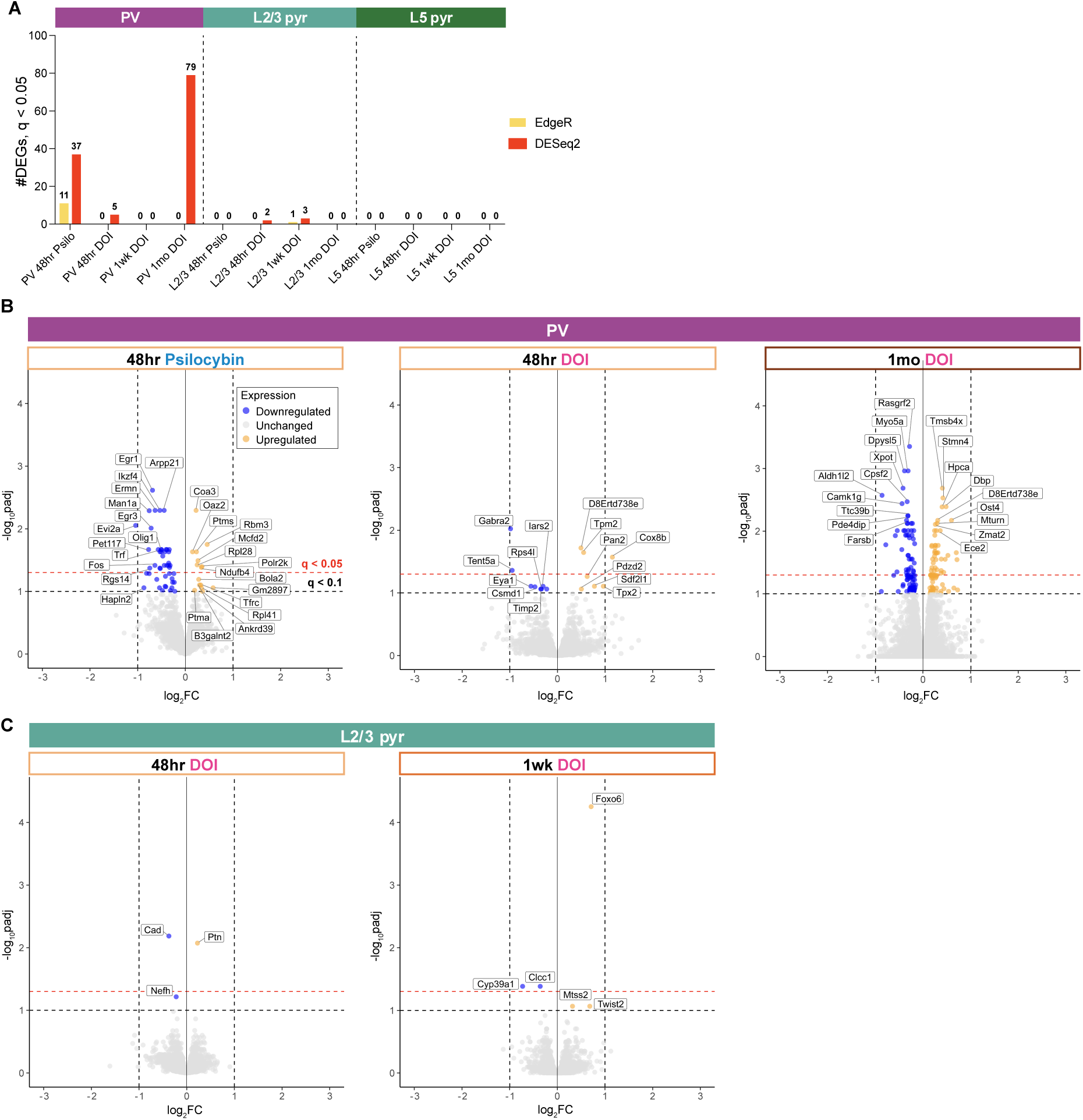
DEGs identified with EdgeR and DESeq2, related to Figure 3. **(A)** Summary of the number of DEGs identified with EdgeR and DESeq2 pipelines, with p_adj_ cutoff of 0.05. **(B)** Volcano plots of DEGs in PV interneurons, 48-hours after DOI or psilocybin, and 1-month after DOI. Cutoffs are for p_adj_ of 0.1 and 0.05. **(C)** Volcano plots of DEGs in L2/3 pyramidal neurons, 48-hours and 1-week after DOI.

**Figure S3.**
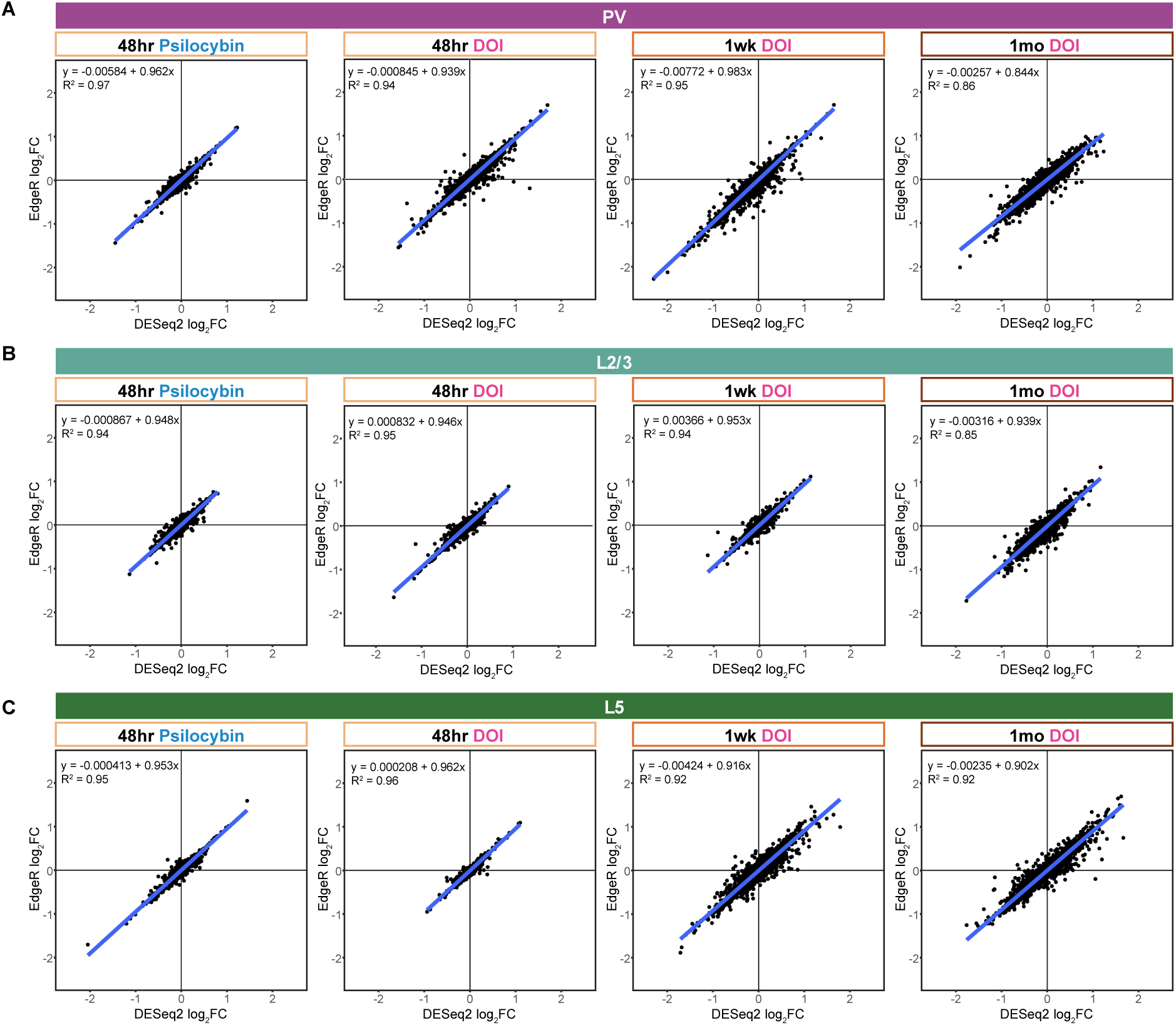
Comparison of EdgeR and DESeq2 for differential gene expression analysis, related to Figure 3. **(A-C)** Scatter plots showing correlation of log_2_FC values from EdgeR and DESeq2 of all genes across (A) PV, (B) L2/3, (C) and L5 treatments/time points.

**Figure S4.**
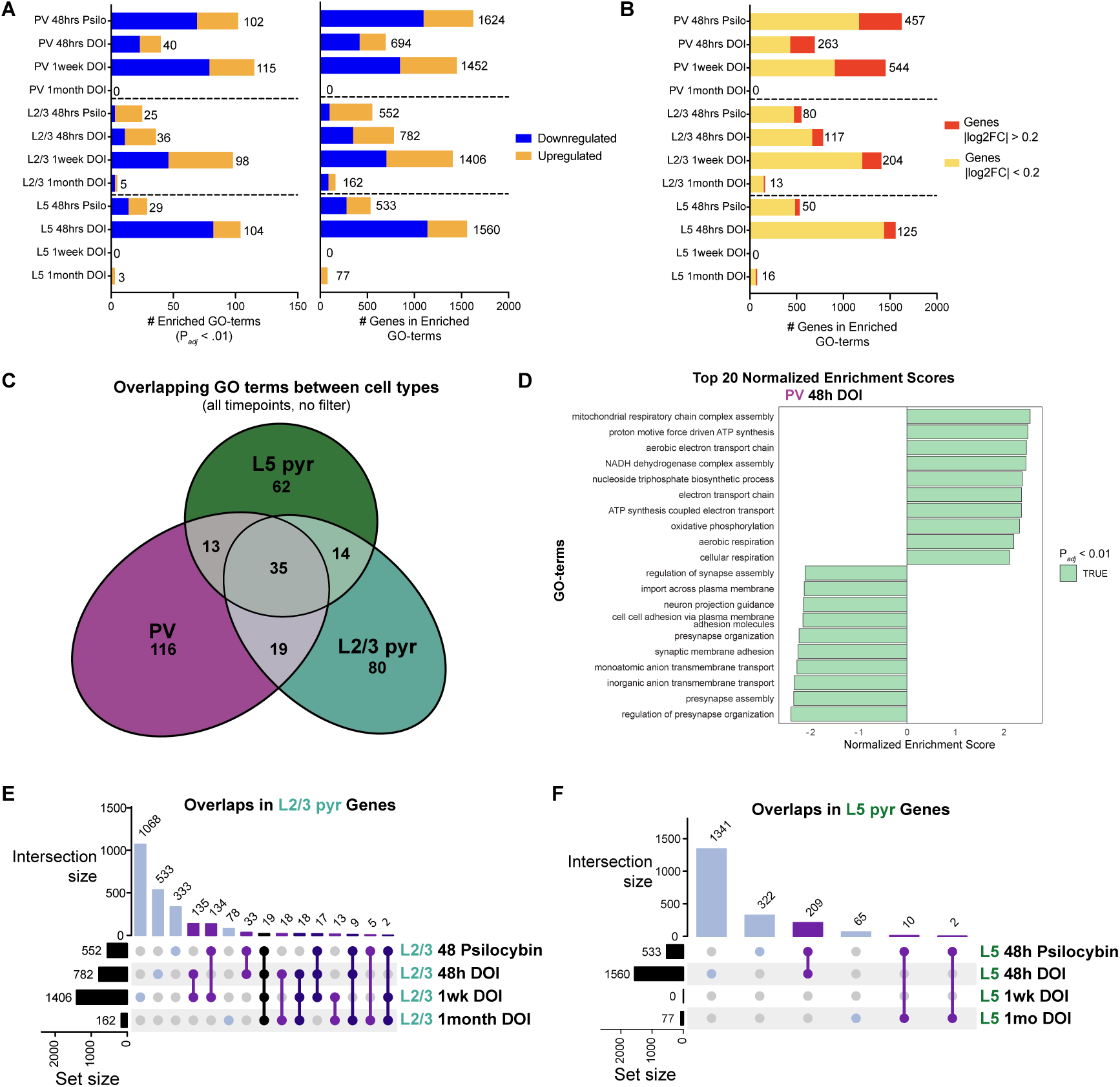
GSEA identified cell type-specific gene expression changes across time points, related to Figure 3. **(A)** Number of enriched GO terms, separated by terms that were up-or down-regulated, and number of genes in enriched GO terms. **(B)** Number of genes in enriched GO terms, separated by genes with |log2FC| > 0.2 or < 0.2. **(C)** Venn diagram showing overlaps in enriched GO terms between cell types, from all conditions. **(D)** Top 10 up– and down-regulated GO terms by normalized enrichment scores (NES) in PV interneurons, 48-hours after DOI. **(E-F)** UpSet plot showing unique and overlapping genes in enriched GO terms between conditions in (E) L2/3 and (F) L5 pyramidal neurons.

**Figure S5.**
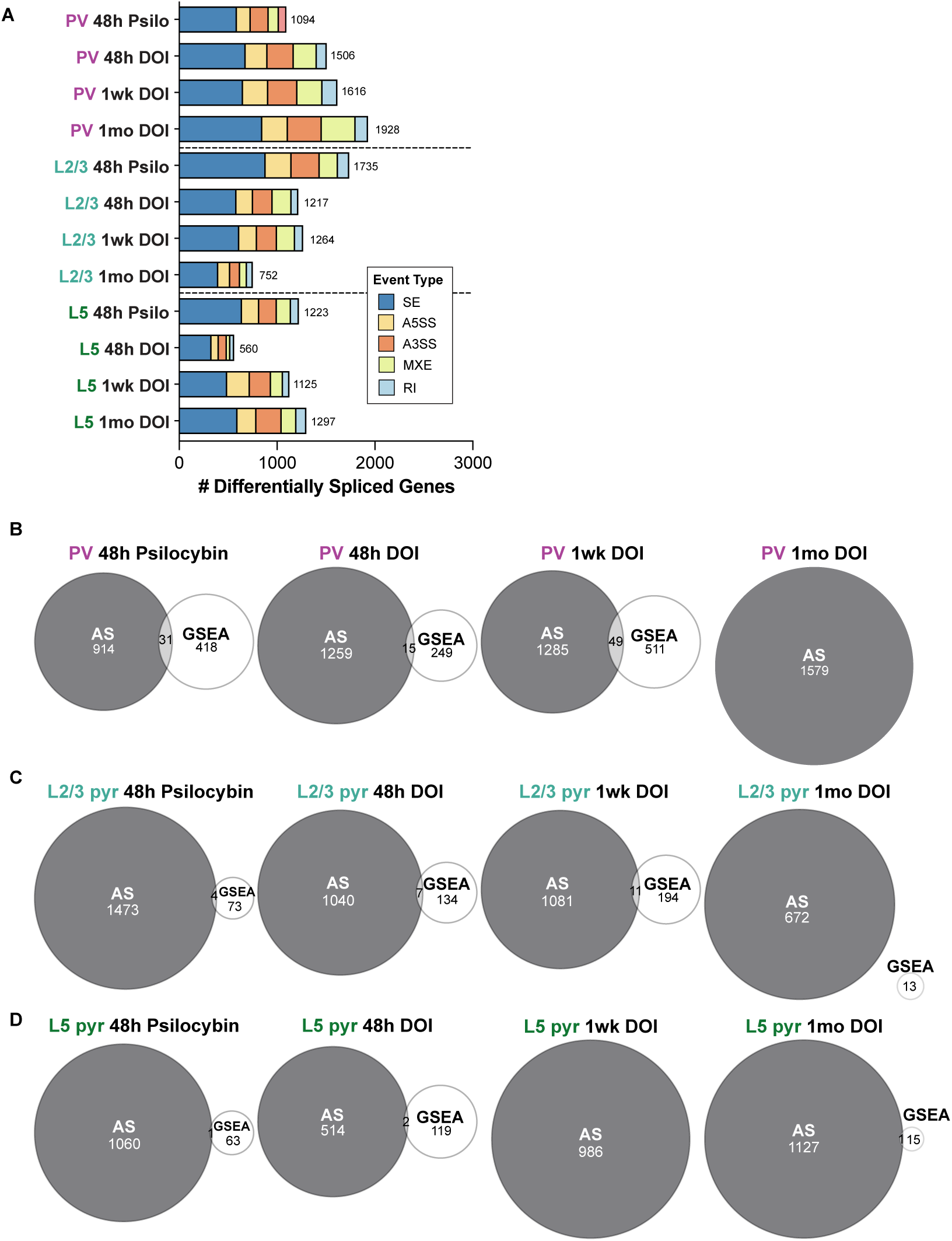
Genes with differentially splicing events identified from alternative splicing analysis, related to Figure 4. **(A)** Number of genes with one or more differential splicing events, separated by event type. **(B-D)** Venn diagram showing overlap of genes identified from alternative splicing analysis vs GSEA, for all conditions.

**Figure S6.**
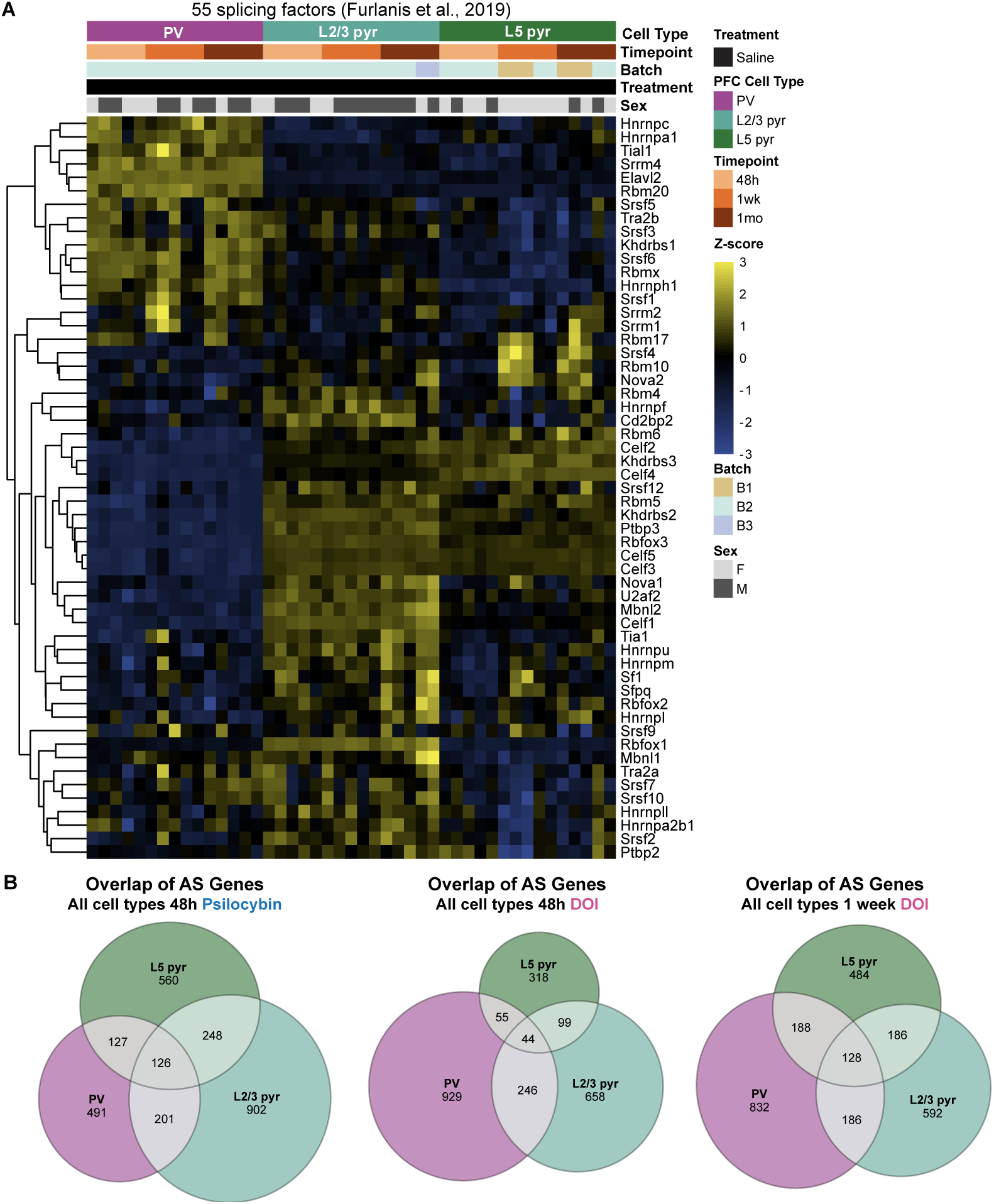
Cell type-specific regulation of alternative splicing, related to Figure 4. **(A)** Heatmap showing gene expression values of bona fide splicing factors from Furlanis et al., 2019 in all saline samples in the current study. **(B)** Venn diagram showing overlap of differentially spliced genes between cell types, for all conditions.

## References

1. Goodwin, G. M., Aaronson, S. T., Alvarez, O., Arden, P. C., Baker, A., Bennett, J. C., Bird, C., Blom, R. E., Brennan, C., Brusch, D., Burke, L., Campbell-Coker, K., Carhart-Harris, R., Cattell, J., Daniel, A., DeBattista, C., Dunlop, B. W., Eisen, K., Feifel, D., … Malievskaia, E. (2022). Single-Dose Psilocybin for a Treatment-Resistant Episode of Major Depression. New England Journal of Medicine, 387(18), 1637–1648. 10.1056/NEJMoa2206443

2. Raison, C. L., Sanacora, G., Woolley, J., Heinzerling, K., Dunlop, B. W., Brown, R. T., Kakar, R., Hassman, M., Trivedi, R. P., Robison, R., Gukasyan, N., Nayak, S. M., Hu, X., O’Donnell, K. C., Kelmendi, B., Sloshower, J., Penn, A. D., Bradley, E., Kelly, D. F., … Griffiths, R. R. (2023). Single-Dose Psilocybin Treatment for Major Depressive Disorder: A Randomized Clinical Trial. JAMA, 330(9), 843. 10.1001/jama.2023.14530

3. Ross, S., Bossis, A., Guss, J., Agin-Liebes, G., Malone, T., Cohen, B., Mennenga, S. E., Belser, A., Kalliontzi, K., Babb, J., Su, Z., Corby, P., & Schmidt, B. L. (2016). Rapid and sustained symptom reduction following psilocybin treatment for anxiety and depression in patients with life-threatening cancer: A randomized controlled trial. Journal of Psychopharmacology, 30(12), 1165–1180. 10.1177/0269881116675512

4. Von Rotz, R., Schindowski, E. M., Jungwirth, J., Schuldt, A., Rieser, N. M., Zahoranszky, K., Seifritz, E., Nowak, A., Nowak, P., Jäncke, L., Preller, K. H., & Vollenweider, F. X. (2023). Single-dose psilocybin-assisted therapy in major depressive disorder: A placebo-controlled, double-blind, randomised clinical trial. eClinicalMedicine, 56, 101809. 10.1016/j.eclinm.2022.101809

5. de la Fuente Revenga, M., Zhu, B., Guevara, C. A., Naler, L. B., Saunders, J. M., Zhou, Z., Toneatti, R., Sierra, S., Wolstenholme, J. T., Beardsley, P. M., Huntley, G. W., Lu, C., & González-Maeso, J. (2021). Prolonged epigenomic and synaptic plasticity alterations following single exposure to a psychedelic in mice. Cell Reports, 37(3), 109836. 10.1016/j.celrep.2021.109836

6. Huang, Z., Wei, X., Wang, Y., Tian, J., Dong, J., Liang, B., Lu, L., & Zhang, W. (2024). Psilocybin Promotes Cell-Type-Specific Changes in the Orbitofrontal Cortex Revealed by Single-Nucleus RNA-seq. 10.1101/2024.01.07.573163

7. Liao, C., O’Farrell, E., Qalieh, Y., Savalia, N. K., Girgenti, M. J., Kwan, K. Y., & Kwan, A. C. (2025). Single-nucleus transcriptomics reveals time-dependent and cell-type-specific effects of psilocybin on gene expression. 10.1101/2025.01.04.631335

8. Muir, J., Lin, S., Aarrestad, I. K., Daniels, H. R., Ma, J., Tian, L., Olson, D. E., & Kim, C. K. (2024). Isolation of psychedelic-responsive neurons underlying anxiolytic behavioral states. Science, 386(6723), 802–810. 10.1126/science.adl0666

9. Nardou, R., Sawyer, E., Song, Y. J., Wilkinson, M., Padovan-Hernandez, Y., De Deus, J. L., Wright, N., Lama, C., Faltin, S., Goff, L. A., Stein-O’Brien, G. L., & Dölen, G. (2023). Psychedelics reopen the social reward learning critical period. Nature, 618(7966), 790– 798. 10.1038/s41586-023-06204-3

10. Zhou, D., Schuler, H., Cvetkovska, V., Meccia, J., Harutyunyan, A. S., Ragoussis, J., & Bagot, R. C. (2025). Single-cell sequencing reveals psilocybin induces sustained cell-type specific plasticity in mouse medial prefrontal cortex. 10.1101/2025.01.08.631940

11. González-Maeso, J., Yuen, T., Ebersole, B. J., Wurmbach, E., Lira, A., Zhou, M., Weisstaub, N., Hen, R., Gingrich, J. A., & Sealfon, S. C. (2003). Transcriptome Fingerprints Distinguish Hallucinogenic and Nonhallucinogenic 5-Hydroxytryptamine 2A Receptor Agonist Effects in Mouse Somatosensory Cortex. The Journal of Neuroscience, 23(26), 8836–8843. 10.1523/JNEUROSCI.23-26-08836.2003

12. Nardou, R., Lewis, E. M., Rothhaas, R., Xu, R., Yang, A., Boyden, E., & Dölen, G. (2019). Oxytocin-dependent reopening of a social reward learning critical period with MDMA. Nature, 569(7754), 116–120. 10.1038/s41586-019-1075-9

13. Pędzich, B. D., Rubens, S., Sekssaoui, M., Pierre, A., Van Schuerbeek, A., Marin, P., Bockaert, J., Valjent, E., Bécamel, C., & De Bundel, D. (2022). Effects of a psychedelic 5-HT2A receptor agonist on anxiety-related behavior and fear processing in mice. Neuropsychopharmacology, 47(7), 1304–1314. 10.1038/s41386-022-01324-2

14. Tiwari, P., Davoudian, P. A., Kapri, D., Vuruputuri, R. M., Karaba, L. A., Sharma, M., Zanni, G., Balakrishnan, A., Chaudhari, P. R., Pradhan, A., Suryavanshi, S., Bath, K. G., Ansorge, M. S., Fernandez-Ruiz, A., Kwan, A. C., & Vaidya, V. A. (2024). Ventral hippocampal parvalbumin interneurons gate the acute anxiolytic action of the serotonergic psychedelic DOI. *Neuron*, S0896627324006408. 10.1016/j.neuron.2024.08.016

15. Shaw, E., & Woolley, D. W. (1956). Some Serotoninlike Activities of Lysergic Acid Diethylamide. Science, 124(3212), 121–122. 10.1126/science.124.3212.121

16. Vollenweider, F. X., & Preller, K. H. (2020). Psychedelic drugs: Neurobiology and potential for treatment of psychiatric disorders. Nature Reviews Neuroscience, 21(11), 611–624. 10.1038/s41583-020-0367-2

17. Woolley, D. W., & Shaw, E. (1954). A BIOCHEMICAL AND PHARMACOLOGICAL SUGGESTION ABOUT CERTAIN MENTAL DISORDERS. Proceedings of the National Academy of Sciences, 40(4), 228–231. 10.1073/pnas.40.4.228

18. Kwan, A. C., Olson, D. E., Preller, K. H., & Roth, B. L. (2022). The neural basis of psychedelic action. Nature Neuroscience, 25(11), 1407–1419. 10.1038/s41593-022-01177-4

19. Fantegrossi, W. E., Simoneau, J., Cohen, M. S., Zimmerman, S. M., Henson, C. M., Rice, K. C., & Woods, J. H. (2010). Interaction of 5-HT2A and 5-HT2C Receptors in R(−)-2,5-Dimethoxy-4-iodoamphetamine-Elicited Head Twitch Behavior in Mice. The Journal of Pharmacology and Experimental Therapeutics, 335(3), 728–734. 10.1124/jpet.110.172247

20. González-Maeso, J., Weisstaub, N. V., Zhou, M., Chan, P., Ivic, L., Ang, R., Lira, A., Bradley-Moore, M., Ge, Y., Zhou, Q., Sealfon, S. C., & Gingrich, J. A. (2007). Hallucinogens Recruit Specific Cortical 5-HT2A Receptor-Mediated Signaling Pathways to Affect Behavior. Neuron, 53(3), 439–452. 10.1016/j.neuron.2007.01.008

21. Madsen, M. K., Fisher, P. M., Burmester, D., Dyssegaard, A., Stenbæk, D. S., Kristiansen, S., Johansen, S. S., Lehel, S., Linnet, K., Svarer, C., Erritzoe, D., Ozenne, B., & Knudsen, G. M. (2019). Psychedelic effects of psilocybin correlate with serotonin 2A receptor occupancy and plasma psilocin levels. Neuropsychopharmacology, 44(7), 1328–1334. 10.1038/s41386-019-0324-9

22. McClure-Begley, T. D., & Roth, B. L. (2022). The promises and perils of psychedelic pharmacology for psychiatry. Nature Reviews Drug Discovery, 21(6), 463–473. 10.1038/s41573-022-00421-7

23. Ray, T. S. (2010). Psychedelics and the Human Receptorome. PLoS ONE, 5(2), e9019. 10.1371/journal.pone.0009019

24. Vollenweider, F. X., Vollenweider-Scherpenhuyzen, M. F. I., Bäbler, A., Vogel, H., & Hell, D. (1998). Psilocybin induces schizophrenia-like psychosis in humans via a serotonin-2 agonist action: NeuroReport, 9(17), 3897–3902. 10.1097/00001756-199812010-00024

25. Pazos, A., Probst, A., & Palacios, J. M. (1987). Serotonin receptors in the human brain—IV. Autoradiographic mapping of serotonin-2 receptors. Neuroscience, 21(1), 123–139. 10.1016/0306-4522(87)90327-7

26. Weber. (2010). Htr2a gene and 5-HT2A receptor expression in the cerebral cortex studied using genetically modified mice. Frontiers in Neuroscience. 10.3389/fnins.2010.00036

27. Etkin, A., & Wager, T. D. (2007). Functional Neuroimaging of Anxiety: A Meta-Analysis of Emotional Processing in PTSD, Social Anxiety Disorder, and Specific Phobia. American Journal of Psychiatry, 164(10), 1476–1488. 10.1176/appi.ajp.2007.07030504

28. Glantz, L. A., & Lewis, D. A. (2000). Decreased Dendritic Spine Density on Prefrontal Cortical Pyramidal Neurons in Schizophrenia. Archives of General Psychiatry, 57(1), 65. 10.1001/archpsyc.57.1.65

29. Nashed, M. G., Seidlitz, E. P., Frey, B. N., & Singh, G. (2015). Depressive-like behaviours and decreased dendritic branching in the medial prefrontal cortex of mice with tumors: A novel validated model of cancer-induced depression. Behavioural Brain Research, 294, 25–35. 10.1016/j.bbr.2015.07.040

30. Radley, J. J., Rocher, A. B., Rodriguez, A., Ehlenberger, D. B., Dammann, M., McEwen, B. S., Morrison, J. H., Wearne, S. L., & Hof, P. R. (2008). Repeated stress alters dendritic spine morphology in the rat medial prefrontal cortex. Journal of Comparative Neurology, 507(1), 1141–1150. 10.1002/cne.21588

31. Xu, P., Chen, A., Li, Y., Xing, X., & Lu, H. (2019). Medial prefrontal cortex in neurological diseases. Physiological Genomics, 51(9), 432–442. 10.1152/physiolgenomics.00006.2019

32. Shao, L.-X., Liao, C., Gregg, I., Davoudian, P. A., Savalia, N. K., Delagarza, K., & Kwan, A. C. (2021). Psilocybin induces rapid and persistent growth of dendritic spines in frontal cortex in vivo. Neuron, 109(16), 2535–2544.e4. 10.1016/j.neuron.2021.06.008

33. Donato, F., Rompani, S. B., & Caroni, P. (2013). Parvalbumin-expressing basket-cell network plasticity induced by experience regulates adult learning. Nature, 504(7479), 272–276. 10.1038/nature12866

34. Kuhlman, S. J., Olivas, N. D., Tring, E., Ikrar, T., Xu, X., & Trachtenberg, J. T. (2013). A disinhibitory microcircuit initiates critical-period plasticity in the visual cortex. Nature, 501(7468), Article 7468. 10.1038/nature12485

35. McFarlan, A. R., Chou, C. Y. C., Watanabe, A., Cherepacha, N., Haddad, M., Owens, H., & Sjöström, P. J. (2023). The plasticitome of cortical interneurons. Nature Reviews Neuroscience, 24(2), Article 2. 10.1038/s41583-022-00663-9

36. Sale, A., Berardi, N., Spolidoro, M., Baroncelli, L., & Maffei, L. (2010). GABAergic inhibition in visual cortical plasticity. Frontiers in Cellular Neuroscience, 4, 10. 10.3389/fncel.2010.00010

37. Cameron, L. P., Patel, S. D., Vargas, M. V., Barragan, E. V., Saeger, H. N., Warren, H. T., Chow, W. L., Gray, J. A., & Olson, D. E. (2023). 5-HT2ARs Mediate Therapeutic Behavioral Effects of Psychedelic Tryptamines. ACS Chemical Neuroscience, 14(3), 351–358. 10.1021/acschemneuro.2c00718

38. Ly, C., Greb, A. C., Cameron, L. P., Wong, J. M., Barragan, E. V., Wilson, P. C., Burbach, K. F., Soltanzadeh Zarandi, S., Sood, A., Paddy, M. R., Duim, W. C., Dennis, M. Y., McAllister, A. K., Ori-McKenney, K. M., Gray, J. A., & Olson, D. E. (2018). Psychedelics Promote Structural and Functional Neural Plasticity. Cell Reports, 23(11), 3170–3182. 10.1016/j.celrep.2018.05.022

39. Mauger, O., Lemoine, F., & Scheiffele, P. (2016). Targeted Intron Retention and Excision for Rapid Gene Regulation in Response to Neuronal Activity. Neuron, 92(6), 1266–1278. 10.1016/j.neuron.2016.11.032

40. Yap, E.-L., & Greenberg, M. E. (2018). Activity-Regulated Transcription: Bridging the Gap between Neural Activity and Behavior. Neuron, 100(2), 330–348. 10.1016/j.neuron.2018.10.013

41. Raj, B., & Blencowe, B. J. (2015). Alternative Splicing in the Mammalian Nervous System: Recent Insights into Mechanisms and Functional Roles. Neuron, 87(1), 14–27. 10.1016/j.neuron.2015.05.004

42. Barbosa-Morais, N. L., Irimia, M., Pan, Q., Xiong, H. Y., Gueroussov, S., Lee, L. J., Slobodeniuc, V., Kutter, C., Watt, S., Çolak, R., Kim, T., Misquitta-Ali, C. M., Wilson, M. D., Kim, P. M., Odom, D. T., Frey, B. J., & Blencowe, B. J. (2012). The Evolutionary Landscape of Alternative Splicing in Vertebrate Species. Science, 338(6114), 1587– 1593. 10.1126/science.1230612

43. Wright, C. J., Smith, C. W. J., & Jiggins, C. D. (2022). Alternative splicing as a source of phenotypic diversity. Nature Reviews Genetics, 23(11), 697–710. 10.1038/s41576-022-00514-4

44. Gomez, A. M., Traunmüller, L., & Scheiffele, P. (2021). Neurexins: Molecular codes for shaping neuronal synapses. Nature Reviews Neuroscience, 22(3), 137–151. 10.1038/s41583-020-00415-7

45. Furlanis, E., Traunmüller, L., Fucile, G., & Scheiffele, P. (2019). Landscape of ribosome-engaged transcript isoforms reveals extensive neuronal-cell-class-specific alternative splicing programs. Nature Neuroscience, 22(10), 1709–1717. 10.1038/s41593-019-0465-5

46. Carvalho, L., & Lasek, A. W. (2024). It is not just about transcription: Involvement of brain RNA splicing in substance use disorders. Journal of Neural Transmission, 131(5), 495– 503. 10.1007/s00702-024-02740-y

47. Feng, J., Wilkinson, M., Liu, X., Purushothaman, I., Ferguson, D., Vialou, V., Maze, I., Shao, N., Kennedy, P., Koo, J., Dias, C., Laitman, B., Stockman, V., LaPlant, Q., Cahill, M. E., Nestler, E. J., & Shen, L. (2014). Chronic cocaine-regulated epigenomic changes in mouse nucleus accumbens. Genome Biology, 15(4), R65. 10.1186/gb-2014-15-4-r65

48. Gandal, M. J., Zhang, P., Hadjimichael, E., Walker, R. L., Chen, C., Liu, S., Won, H., van Bakel, H., Varghese, M., Wang, Y., Shieh, A. W., Haney, J., Parhami, S., Belmont, J., Kim, M., Moran Losada, P., Khan, Z., Mleczko, J., Xia, Y., … Geschwind, D. H. (2018). Transcriptome-wide isoform-level dysregulation in ASD, schizophrenia, and bipolar disorder. Science, 362(6420), eaat8127. 10.1126/science.aat8127

49. Traunmüller, L., Gomez, A. M., Nguyen, T.-M., & Scheiffele, P. (2016). Control of neuronal synapse specification by a highly dedicated alternative splicing program. Science, 352(6288), 982–986. 10.1126/science.aaf2397

50. Bhattacherjee, A., Djekidel, M. N., Chen, R., Chen, W., Tuesta, L. M., & Zhang, Y. (2019). Cell type-specific transcriptional programs in mouse prefrontal cortex during adolescence and addiction. Nature Communications, 10(1), 4169. 10.1038/s41467-019-12054-3

51. Bhattacherjee, A., Zhang, C., Watson, B. R., Djekidel, M. N., Moffitt, J. R., & Zhang, Y. (2023). Spatial transcriptomics reveals the distinct organization of mouse prefrontal cortex and neuronal subtypes regulating chronic pain. Nature Neuroscience, 26(11), 1880–1893. 10.1038/s41593-023-01455-9

52. Sanz, E., Yang, L., Su, T., Morris, D. R., McKnight, G. S., & Amieux, P. S. (2009). Cell-type-specific isolation of ribosome-associated mRNA from complex tissues. Proceedings of the National Academy of Sciences, 106(33), 13939–13944. 10.1073/pnas.0907143106

53. Blevins, W. R., Tavella, T., Moro, S. G., Blasco-Moreno, B., Closa-Mosquera, A., Díez, J., Carey, L. B., & Albà, M. M. (2019). Extensive post-transcriptional buffering of gene expression in the response to severe oxidative stress in baker’s yeast. Scientific Reports, 9(1), 11005. 10.1038/s41598-019-47424-w

54. Glock, C., Biever, A., Tushev, G., Nassim-Assir, B., Kao, A., Bartnik, I., Tom Dieck, S., & Schuman, E. M. (2021). The translatome of neuronal cell bodies, dendrites, and axons. Proceedings of the National Academy of Sciences, 118(43), e2113929118. 10.1073/pnas.2113929118

55. Harris, J. A., Hirokawa, K. E., Sorensen, S. A., Gu, H., Mills, M., Ng, L. L., Bohn, P., Mortrud, M., Ouellette, B., Kidney, J., Smith, K. A., Dang, C., Sunkin, S., Bernard, A., Oh, S. W., Madisen, L., & Zeng, H. (2014). Anatomical characterization of Cre driver mice for neural circuit mapping and manipulation. Frontiers in Neural Circuits, 8. 10.3389/fncir.2014.00076

56. Hrvatin, S., Hochbaum, D. R., Nagy, M. A., Cicconet, M., Robertson, K., Cheadle, L., Zilionis, R., Ratner, A., Borges-Monroy, R., Klein, A. M., Sabatini, B. L., & Greenberg, M. E. (2018). Single-cell analysis of experience-dependent transcriptomic states in the mouse visual cortex. Nature Neuroscience, 21(1), 120–129. 10.1038/s41593-017-0029-5

57. Wang, X., Park, J., Susztak, K., Zhang, N. R., & Li, M. (2019). Bulk tissue cell type deconvolution with multi-subject single-cell expression reference. Nature Communications, 10(1), 380. 10.1038/s41467-018-08023-x

58. Subramanian, A., Tamayo, P., Mootha, V. K., Mukherjee, S., Ebert, B. L., Gillette, M. A., Paulovich, A., Pomeroy, S. L., Golub, T. R., Lander, E. S., & Mesirov, J. P. (2005). Gene set enrichment analysis: A knowledge-based approach for interpreting genome-wide expression profiles. Proceedings of the National Academy of Sciences, 102(43), 15545– 15550. 10.1073/pnas.0506580102

59. Aboharb, F., Davoudian, P. A., Shao, L.-X., Liao, C., Rzepka, G. N., Wojtasiewicz, C., Indajang, J., Dibbs, M., Rondeau, J., Sherwood, A. M., Kaye, A. P., & Kwan, A. C. (2024). Classification of psychedelics and psychoactive drugs based on brain-wide imaging of cellular c-Fos expression. Neuroscience. 10.1101/2024.05.23.590306

60. Davoudian, P. A., Shao, L.-X., & Kwan, A. C. (2023). Shared and Distinct Brain Regions Targeted for Immediate Early Gene Expression by Ketamine and Psilocybin. ACS Chemical Neuroscience, 14(3), 468–480. 10.1021/acschemneuro.2c00637

61. Rijsketic, D. R., Casey, A. B., Barbosa, D. A. N., Zhang, X., Hietamies, T. M., Ramirez-Ovalle, G., Pomrenze, M. B., Halpern, C. H., Williams, L. M., Malenka, R. C., & Heifets, B. D. (2023). UNRAVELing the synergistic effects of psilocybin and environment on brain-wide immediate early gene expression in mice. Neuropsychopharmacology, 48(12), 1798–1807. 10.1038/s41386-023-01613-4

62. Jacobs, A., & Elmer, K. R. (2021). Alternative splicing and gene expression play contrasting roles in the parallel phenotypic evolution of a salmonid fish. Molecular Ecology, 30(20), 4955–4969. 10.1111/mec.15817

63. Keren, H., Lev-Maor, G., & Ast, G. (2010). Alternative splicing and evolution: Diversification, exon definition and function. Nature Reviews Genetics, 11(5), 345–355. 10.1038/nrg2776

64. Besco, J. A., Frostholm, A., Popesco, M. C., Burghes, A. H., & Rotter, A. (2001). Genomic organization and alternative splicing of the human and mouse RPTPρ genes. BMC Genomics, 2(1), 1. 10.1186/1471-2164-2-1

65. Han, B. Y., Seah, M. K. Y., Brooks, I. R., Quek, D. H. P., Huxley, D. R., Foo, C.-S., Lee, L. T., Wollmann, H., Guo, H., Messerschmidt, D. M., & Guccione, E. (2020). Global translation during early development depends on the essential transcription factor PRDM10. Nature Communications, 11(1), 3603. 10.1038/s41467-020-17304-3

66. Lu, Y. C., Nazarko, O. V., Sando, R., Salzman, G. S., Li, N.-S., Südhof, T. C., & Araç, D. (2015). Structural Basis of Latrophilin-FLRT-UNC5 Interaction in Cell Adhesion. Structure, 23(9), 1678–1691. 10.1016/j.str.2015.06.024

67. Yang, Z., Zhou, D., Li, H., Cai, X., Liu, W., Wang, L., Chang, H., Li, M., & Xiao, X. (2020). The genome-wide risk alleles for psychiatric disorders at 3p21.1 show convergent effects on mRNA expression, cognitive function, and mushroom dendritic spine. Molecular Psychiatry, 25(1), 48–66. 10.1038/s41380-019-0592-0

68. Beurdeley, M., Spatazza, J., Lee, H. H. C., Sugiyama, S., Bernard, C., Di Nardo, A. A., Hensch, T. K., & Prochiantz, A. (2012). Otx2 Binding to Perineuronal Nets Persistently Regulates Plasticity in the Mature Visual Cortex. Journal of Neuroscience, 32(27), 9429– 9437. 10.1523/JNEUROSCI.0394-12.2012

69. Favuzzi, E., Marques-Smith, A., Deogracias, R., Winterflood, C. M., Sánchez-Aguilera, A., Mantoan, L., Maeso, P., Fernandes, C., Ewers, H., & Rico, B. (2017). Activity-Dependent Gating of Parvalbumin Interneuron Function by the Perineuronal Net Protein Brevican. Neuron, 95(3), 639–655.e10. 10.1016/j.neuron.2017.06.028

70. Reh, R. K., Dias, B. G., Nelson, C. A., Kaufer, D., Werker, J. F., Kolb, B., Levine, J. D., & Hensch, T. K. (2020). Critical period regulation across multiple timescales. Proceedings of the National Academy of Sciences, 117(38), 23242–23251. 10.1073/pnas.1820836117

71. Ye, Q., & Miao, Q. (2013). Experience-dependent development of perineuronal nets and chondroitin sulfate proteoglycan receptors in mouse visual cortex. Matrix Biology, 32(6), 352–363. 10.1016/j.matbio.2013.04.001

72. Hensch, T. K. (2005). Critical period plasticity in local cortical circuits. Nature Reviews Neuroscience, 6(11), 877–888. 10.1038/nrn1787

73. Pizzorusso, T., Medini, P., Berardi, N., Chierzi, S., Fawcett, J. W., & Maffei, L. (2002). Reactivation of Ocular Dominance Plasticity in the Adult Visual Cortex. Science, 298(5596), 1248–1251. 10.1126/science.1072699

74. Pizzorusso, T., Medini, P., Landi, S., Baldini, S., Berardi, N., & Maffei, L. (2006). Structural and functional recovery from early monocular deprivation in adult rats. Proceedings of the National Academy of Sciences, 103(22), 8517–8522. 10.1073/pnas.0602657103

75. Brückner, G., Härtig, W., Kacza, J., Seeger, J., Welt, K., & Brauer, K. (1996). Extracellular matrix organization in various regions of rat brain grey matter. Journal of Neurocytology, 25(1), 333–346. 10.1007/BF02284806

76. Enwright, J. F., Sanapala, S., Foglio, A., Berry, R., Fish, K. N., & Lewis, D. A. (2016). Reduced Labeling of Parvalbumin Neurons and Perineuronal Nets in the Dorsolateral Prefrontal Cortex of Subjects with Schizophrenia. Neuropsychopharmacology, 41(9), 2206–2214. 10.1038/npp.2016.24

77. Fawcett, J. W., Oohashi, T., & Pizzorusso, T. (2019). The roles of perineuronal nets and the perinodal extracellular matrix in neuronal function. Nature Reviews Neuroscience, 20(8), 451–465. 10.1038/s41583-019-0196-3

78. Lensjø, K. K., Lepperød, M. E., Dick, G., Hafting, T., & Fyhn, M. (2017). Removal of Perineuronal Nets Unlocks Juvenile Plasticity Through Network Mechanisms of Decreased Inhibition and Increased Gamma Activity. The Journal of Neuroscience, 37(5), 1269–1283. 10.1523/JNEUROSCI.2504-16.2016

79. Lupori, L., Totaro, V., Cornuti, S., Ciampi, L., Carrara, F., Grilli, E., Viglione, A., Tozzi, F., Putignano, E., Mazziotti, R., Amato, G., Gennaro, C., Tognini, P., & Pizzorusso, T. (2023). A comprehensive atlas of perineuronal net distribution and colocalization with parvalbumin in the adult mouse brain. Cell Reports, 42(7), 112788. 10.1016/j.celrep.2023.112788

80. Anastasiades, P. G., & Carter, A. G. (2021). Circuit organization of the rodent medial prefrontal cortex. Trends in Neurosciences, 44(7), 550–563. 10.1016/j.tins.2021.03.006

81. Hu, H., Gan, J., & Jonas, P. (2014). Fast-spiking, parvalbumin+ GABAergic interneurons: From cellular design to microcircuit function. Science, 345(6196), 1255263. 10.1126/science.1255263

82. Rupert, D. D., & Shea, S. D. (2022). Parvalbumin-Positive Interneurons Regulate Cortical Sensory Plasticity in Adulthood and Development Through Shared Mechanisms. Frontiers in Neural Circuits, 16, 886629. 10.3389/fncir.2022.886629

83. Sugiyama, S., Nardo, A. A. D., Aizawa, S., Matsuo, I., Volovitch, M., Prochiantz, A., & Hensch, T. K. (2008). Experience-Dependent Transfer of Otx2 Homeoprotein into the Visual Cortex Activates Postnatal Plasticity. Cell, 134(3), 508–520. 10.1016/j.cell.2008.05.054

84. Wu, Y. K., Miehl, C., & Gjorgjieva, J. (2022). Regulation of circuit organization and function through inhibitory synaptic plasticity. Trends in Neurosciences, 45(12), 884–898. 10.1016/j.tins.2022.10.006

85. Liu, Y., Huang, J., Pandey, R., Liu, P., Therani, B., Qiu, Q., Rao, S., Geurts, A. M., Cowley, A. W., Greene, A. S., & Liang, M. (2023). Robustness of single-cell RNA-seq for identifying differentially expressed genes. BMC Genomics, 24(1), 371. 10.1186/s12864-023-09487-y

86. Fagerholm, E. D., Leech, R., Williams, S., Zarate, C. A., Moran, R. J., & Gilbert, J. R. (2021). Fine-tuning neural excitation/inhibition for tailored ketamine use in treatment-resistant depression. Translational Psychiatry, 11(1), 1–10. 10.1038/s41398-021-01442-3

87. Ferguson, B. R., & Gao, W.-J. (2018). PV Interneurons: Critical Regulators of E/I Balance for Prefrontal Cortex-Dependent Behavior and Psychiatric Disorders. Frontiers in Neural Circuits, 12. 10.3389/fncir.2018.00037

88. Lewis, D. A., Hashimoto, T., & Volk, D. W. (2005). Cortical inhibitory neurons and schizophrenia. Nature Reviews Neuroscience, 6(4), 312–324. 10.1038/nrn1648

89. Yin, Y.-Y., Wang, Y.-H., Liu, W.-G., Yao, J.-Q., Yuan, J., Li, Z.-H., Ran, Y.-H., Zhang, L.-M., & Li, Y.-F. (2021). The role of the excitation:inhibition functional balance in the mPFC in the onset of antidepressants. Neuropharmacology, 191, 108573. 10.1016/j.neuropharm.2021.108573

90. Isaacson, J. S., & Scanziani, M. (2011). How Inhibition Shapes Cortical Activity. Neuron, 72(2), 231–243. 10.1016/j.neuron.2011.09.027

91. Tang, Z.-H., Yu, Z.-P., Li, Q., Zhang, X.-Q., Muhetaer, K., Wang, Z.-C., Xu, P., & Shen, H.-W. (2023). The effects of serotonergic psychedelics in synaptic and intrinsic properties of neurons in layer II/III of the orbitofrontal cortex. Psychopharmacology, 240(6), 1275– 1285. 10.1007/s00213-023-06366-y

92. Lepow, L., Morishita, H., & Yehuda, R. (2021). Critical Period Plasticity as a Framework for Psychedelic-Assisted Psychotherapy. Frontiers in Neuroscience, 15. 10.3389/fnins.2021.710004

93. Miller, M. N., Okaty, B. W., Kato, S., & Nelson, S. B. (2011). Activity-dependent changes in the firing properties of neocortical fast-spiking interneurons in the absence of large changes in gene expression. Developmental Neurobiology, 71(1), 62–70. 10.1002/dneu.20811

94. Hu, Q., Kim, E. J., Feng, J., Grant, G. R., & Heller, E. A. (2017). Histone posttranslational modifications predict specific alternative exon subtypes in mammalian brain. PLOS Computational Biology, 13(6), e1005602. 10.1371/journal.pcbi.1005602

95. Heiman, M., Kulicke, R., Fenster, R. J., Greengard, P., & Heintz, N. (2014). Cell type– specific mRNA purification by translating ribosome affinity purification (TRAP). Nature Protocols, 9(6), 1282–1291. 10.1038/nprot.2014.085

96. Cabrera-Garcia, D. (2022). Patch-clamp data analysis in Python: Passive membrane properties. Spikes and Bursts. https://spikesandbursts.wordpress.com/2022/05/13/patch-clamp-data-analysis-python-passive-membrane-properties/

97. Li, B., & Dewey, C. N. (2011). RSEM: Accurate transcript quantification from RNA-Seq data with or without a reference genome. BMC Bioinformatics, 12(1), 323. 10.1186/1471-2105-12-323

98. Korotkevich, G., Sukhov, V., Budin, N., Shpak, B., Artyomov, M. N., & Sergushichev, A. (2016). Fast gene set enrichment analysis. 10.1101/060012

99. Wang, Y., Xie, Z., Kutschera, E., Adams, J. I., Kadash-Edmondson, K. E., & Xing, Y. (2024). rMATS-turbo: An efficient and flexible computational tool for alternative splicing analysis of large-scale RNA-seq data. Nature Protocols, 19(4), 1083–1104. 10.1038/s41596-023-00944-2

100. Leek, J. T., Johnson, W. E., Parker, H. S., Jaffe, A. E., & Storey, J. D. (2012). The sva package for removing batch effects and other unwanted variation in high-throughput experiments. Bioinformatics, 28(6), 882–883. 10.1093/bioinformatics/bts034

101. Paxinos, G., & Franklin, K. B. J. (2019). Paxinos and Franklin’s the Mouse Brain in Stereotaxic Coordinates. https://shop.elsevier.com/books/paxinos-and-franklins-the-mouse-brain-in-stereotaxic-coordinates/paxinos/978-0-12-816157-9

